# Effective imaging and treatment of Acute Myeloid Leukemia with radiotheranostics targeting the activated conformation of integrin-βeta2

**DOI:** 10.1101/2025.08.21.671206

**Authors:** Anju Wadhwa, Haley Johnson, Kondapa Naidu Bobba, Anil P. Bidkar, Ellis Mayne, Sham Rampersaud, Kamal Mandal, Abhilash Barpanda, Sanjana Prudhvi, Amrik S. Kang, Nancy Greenland, Dana Balitzer, Robin Peter, Athira Raveendran, Shubhankar Naik, Megha Basak, Corynn Kasap, Juwita Huebner, Marina Lopez Alvarez, Sanghee Lee, Veronica Steri, Jarret J. Adams, Sachdev S. Sidhu, David M. Wilson, Youngho Seo, Henry F. VanBrocklin, Aaron C. Logan, Arun P. Wiita, Robert R. Flavell

**Affiliations:** Department of Radiology and Biomedical Imaging, University of California, San Francisco, California 94143, United States; UCSF Helen Diller Family Comprehensive Cancer Center, San Francisco, California; Department of Nuclear Engineering, University of Tennessee-Knoxville, Knoxville, Tennessee 37996, United States; Animal Biotechnology, Gujarat Biotechnology University, Gujarat, Ahmedabad, India; Department of Pathology, University of California, San Francisco, California 94110, United States; University of California, Berkeley, California, United States; Department of Medicine, Division of Hematology/Oncology University of California, San Francisco, California 94110, United States; School of Pharmacy, University of Waterloo, Kitchener, Ontario, Canada N2G 1C5; Department of Laboratory Medicine, University of California, San Francisco, California; Department of Bioengineering and Therapeutic Sciences, University of California, San Francisco, California; Chan Zuckerberg Biohub, San Francisco, California; Department of Pharmaceutical Chemistry, University of California, San Francisco, California 94158-2517, United States

## Abstract

There remains an unmet clinical need for improved treatment strategies in Acute Myeloid Leukemia (AML). Although radiopharmaceutical therapies targeting non-cancer-selective antigens have shown promise in AML, their clinical utility is often limited by prolonged bone marrow suppression. Using a unique proteomics-based strategy, we recently identified the active conformation of integrin-β2 (aITGB2) as a novel, tumor-selective target for AML. Importantly, this conformational epitope is expressed widely on AML cells but minimally on normal marrow progenitors/healthy tissues. Here we first confirmed widespread aITGB2 expression on AML tumors that was largely independent of tumor genotype or prior therapeutic regimen. We developed diagnostic and therapeutic radiopharmaceuticals targeting aITGB2 utilizing a conformation-specific antibody (clone 7065). PET/CT imaging with ^89^Zr and ^134^Ce-labeled 7065 in AML models revealed high target-mediated uptake, greater than that compared to standard of care [^18^F]-FDG. PET/CT imaging with [^89^Zr]DFO*-7065 showed reduced binding to normal bone marrow and immune cells in humanized immune system mice compared to [^89^Zr]DFO*-anti-CD33. For therapy, we developed [^225^Ac]Macropa-PEG_4_-7065 using an optimized chelator-linker combination. Treatment with [^225^Ac]Macropa-PEG_4_-7065 in Nomo-1 and PDX AML disseminated models delayed tumor growth and improved overall survival compared to controls, including [^225^Ac]DOTA-anti-CD33, a clinical stage-radioimmunotherapy under evaluation in AML. Relapsed tumors demonstrated persistent aITGB2 expression, supporting continued development of fractionated dosing schemes, and proteomics analysis indicated activation of TCA cycle and carbon metabolism pathways, consistent with therapy-induced stress responses. These findings highlight [^89^Zr]DFO*-7065 and [^225^Ac]Macropa-7065 as a promising aITGB2-targeted theranostic pair with potential for imaging and treatment in future clinical translation.

**One Sentence Summary:** This study demonstrates promising preclinical efficacy of aITGB2-targeted radiotheranostics for selective imaging and therapy in AML.

## Introduction

Acute myeloid leukemia (AML) is diagnosed in >20,000 Americans per year but has dismal outcomes with <35% 5-year survival (*1*). Despite recent approval of several new small molecule drugs, these agents only lead to modest lifespan extension (*2*). AML is thus widely acknowledged as the greatest unmet medical need among hematologic malignancies. The current approach to AML treatment involves induction therapy with chemotherapy (*3*) (e.g., cytarabine and anthracyclines) to achieve remission, followed by consolidation with high-dose cytarabine or allogeneic stem cell transplantation for eligible patients (*4*). Notably, surface antigen-targeted immunotherapies including monoclonal antibodies and CAR T-cells have shown remarkable clinical efficacy in other blood cancers. However, similar therapeutics in AML thus far remain of limited utility as all leading targets (CD33, CD123, CLL-1) have significant expression on normal hematopoietic and/or endothelial cells, leading to “on target, off tumor” toxicity concerns (*5–6*).

Radiopharmaceutical therapy is a rapidly growing treatment strategy employing cancer-selective targeting vectors to deliver high doses of radiation to cancer cells while sparing healthy tissues (*7*). Interest in this treatment modality has been reinvigorated through recent FDA approvals of ^177^Lu-DOTATATE (Lutathera®) (*8*) and ^177^Lu-PSMA-617 (Pluvicto®) (*9*) which are now standard treatments for neuroendocrine tumors and prostate cancer. When the targeting vector for radiopharmaceutical therapy is an antibody, it is termed radioimmunotherapy (RIT), a promising treatment modality in hematologic malignancies (*10*). In AML, prior efforts using RIT have demonstrated efficacy but also considerable toxicity. A CD33 targeting RIT utilizing [^225^Ac]DOTA-lintuzumab as a single agent (*11*) or in combination with chemotherapeutic against Venetoclax (*12*), CLAG-M (*13,14*) (cladribine; cytarabine; G-CSF; Mitoxantrone), or ASRTX-727 (*15*), showed clinical activity in relapsed/refractory patients with adverse cytogenetics or those who have undergone multiple prior treatments. However, patients experienced hematologic adverse events, such as prolonged neutropenia, infections, and hepatic toxicity (*11*). The CD33 antigen is expressed widely on normal myeloid cells and has only moderate density on tumors (averaging 10,000 CD33 molecule per AML blast) leading to an unfavorable therapeutic index (*16–18*). In addition, anti-CD45 RIT including [^90^Y]-labelled ([^90^Y]-anti-BC8) and [^131^I]-labeled [^131^I]-BC8 have been effective for pre-transplant conditioning by ablating both leukemic and normal hematopoietic cells but by design lead to severe hematologic toxicity, restricting use to fully myeloablative applications (*19,20*). Although these RITs highlight the potential of targeted radiotherapy in AML, challenges such as “on-target, off-tumor” toxicity, myelosuppression, and antigen specificity remain significant barriers. Therefore, despite the high demand for novel targeted therapies, no RIT has been become standard of care (*18*). Advances in antigen selection, tumor-specific targeting, and improved isotope delivery are critical for enhancing the safety and efficacy of RIT for AML treatment.

Through a unique proteomic-based strategy, we recently discovered the active conformation of integrin-b2 (aITGB2) as a novel, tumor-selective immunotherapy target for AML (*21*). We developed an antibody (**clone 7065**) that recognized, in a conformation-selective manner, aITGB2 but not total ITGB2. We showed that a fragment derived from 7065 could be incorporated into highly efficacious chimeric antigen receptor (CAR) T-cells with minimal toxicity to normal hematopoietic tissue. The phenotype validated similar efficacy, but significant safety improvement, for AML immunotherapy of 7065-based CAR-Ts when compared to clinically evaluated CD33-targeting CAR-Ts. However, CAR-T cells in AML have thus far only shown modest clinical efficacy, potentially due to a highly immunosuppressive tumor microenvironment (*22*) or cytokine-mediated growth of tumor cells stimulated by CAR-T activity (*23*). The discovery of our novel target and proprietary conformation-selective 7065 antibody clone thus motivated us to develop additional therapeutic modalities that do not depend on the tumor microenvironment for efficacy.

Herein, we first further validate aITGB2 as a target in AML. We confirm consistent aITGB2 expression across primary bulk tumor cells as well as leukemic stem cells and confirm equivalent antigen expression in both newly diagnosed and relapsed tumors. Leveraging these results, we moved toward the development and evaluation of aITGB2-directed theranostics in AML. [^89^Zr]DFO*-7065 ImmunoPET imaging was evaluated in AML disseminated murine models, which demonstrated targeted mediated imaging with high tumoral uptake, improved compared to the standard of care [^18^F]-FDG. We performed radioimmunotherapy with [^225^Ac]Macropa-PEG_4_-7065 in disseminated cell line and patient-derived xenograft models of AML, with a marked reduction in tumor burden and improvement in overall survival. Importantly, we observed significantly improved performance compared to a previously described anti-CD33 RIT, while we simultaneously found minimal evidence of marrow suppression in murine models. In cases of relapse, tumor proteomic analysis pointed to alterations in adhesion and internalization. Taken together, these results demonstrated the efficacy of aITGB2 targeting theranostics in AML and strongly support future clinical translation.

## Results

### aITGB2 is widely expressed and independent of other AML molecular features

For validation of aITGB2 as an AML specific therapeutic target, we performed flow cytometry to measure the expression of aITGB2 on AML cell lines, demonstrating heterogeneous but generally high aITGB2 expression (Nomo-1 ≥ HL-60 > THP-1 > MV411) (**Figure 1A**). All cell lines had greater aITGB2 expression than a Nomo-1 ITGB2-KO cell line where *ITGB2* was knocked out by CRISPR/Cas9 (**Figure 1A**). We used flow cytometry to examine expression of aITGB2 in 15 primary AML patient samples collected at the time of initial clinical diagnosis, which included a range of common AML genetic abnormalities. These include mutations in *FLT3* (n=10), *DNMT3A* (n=7) samples, *NPM1* (n=5 samples), and *TET2* (n=5), and chromosomal abnormalities +8 (n=6), +5 (n=3), *KMT2A*/*MLL* rearrangement (n=2), -7 (n=2), and -5 (n=1) (**Table S1, S2**). Consistent with our prior work (*21*) we found that 11/15 (73%) primary samples had a median fluorescence intensity (MFI) for aITGB2 at least 2-fold greater than a staining negative control, which we designated as the cutoff for positive expression of aITGB2 (**Figure 1B**). Flow cytometry of Lin-/CD34+/CD38-tumor fraction enriched for leukemic stem cells also showed similar levels of aITGB2 expression to bulk blasts (**Figure S1**). As an aITGB2-targeting therapy would likely be evaluated for clinical translation in a relapsed and/or refractory AML patient population, frequently with prior azacytidine and/or venetoclax treatment, we sought to characterize aITGB2 expression in this population as well. We used flow cytometry to evaluate the percent expression and median fluorescence intensity in matched AML samples from 7 additional patients, taken at initial diagnosis and following refractory disease or relapse (r/r) after at least one round of azacitidine and/or venetoclax (**Table S3**). Notably, we saw no mean decrease in percent expression of aITGB2 and a moderate, albeit non-statistically significant, 1.4-fold increase in median fluorescence intensity in r/r samples compared to those collected at diagnosis (**Figure 1C**). No significant differences were seen in aITGB2 expression between the major genetic subtypes of AML between the primary samples evaluated, suggesting the expression of this target is orthogonal to current molecularly based AML classifications, and therefore not limited to a specific tumor genotype for efficacy.

**Figure 1.**
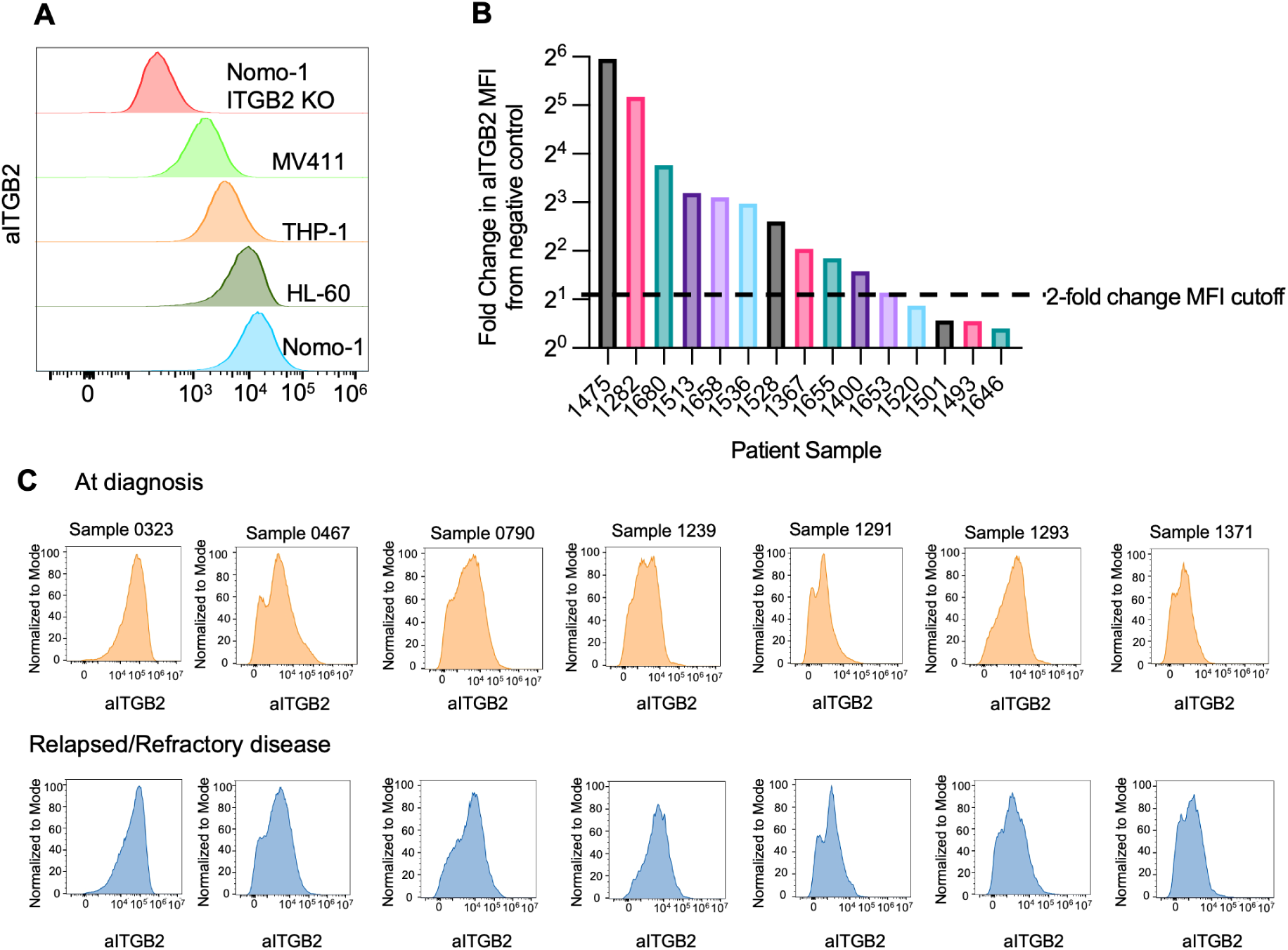
aITGB2 is highly expressed in cell lines and human AML samples. A) Flow Cytometry analysis of aITGB2 in AML cell lines B) Cell surface expression of aITGB2 in AML patient samples C) Comparison of cell surface expression of aITGB2 in patient leukemia samples at time of initial diagnosis and relapse.

### Design and synthesis of radioconjugates targeting aITGB2

IgG 7065 was selected for imaging and therapy due to its high selectivity for aITGB2 and minimal off-target binding (*21*). For imaging, DFO* was used as the chelator for its strong 89Zr binding and reduced non-specific bone uptake—critical for studying hematologic malignancies in bone-localized models (*24–25*). We synthesized four radioimmunoconjugates for imaging and three for therapy **(Figure 2)**. DFO*-7065, DFO*-anti-CD33, and DFO*-IgG were prepared via DFO*-NCS conjugation to 7065, Lintuzumab, and non-targeting IgG, with chelator-to-antibody ratios between 0.4-1.5 **(Figure 2A–C, S2A–C)**. Radiolabeling with Zr-oxalate yielded [^89^Zr]DFO*-7065 at 80.2 ± 0.19% (n=6), with 5.75 mCi/mg specific activity **(Figure S3A–B)**. Size exclusion chromatography confirmed no aggregation post-labeling of [^89^Zr]DFO*-7065 **(Figure S4A–B)**. [^89^Zr]DFO*-anti-CD33 and [^89^Zr]DFO*-IgG were obtained at 82.2 ± 0.58% (n=2) and 86.2 ± 0.34% (n=5), with specific activities of 4.37 and 7.5 mCi/mg **(Figure S5A–D)**. All radio conjugates demonstrated purity >98%. A LINDMO assay in Nomo-1 cells showed [^89^Zr]DFO*-7065 had 87.8 ± 0.65% immunoreactivity **(Figure S6A)**, and >99% stability in saline and human serum over seven days at 37°C **(Figure S6B)**.

**Figure 2.**
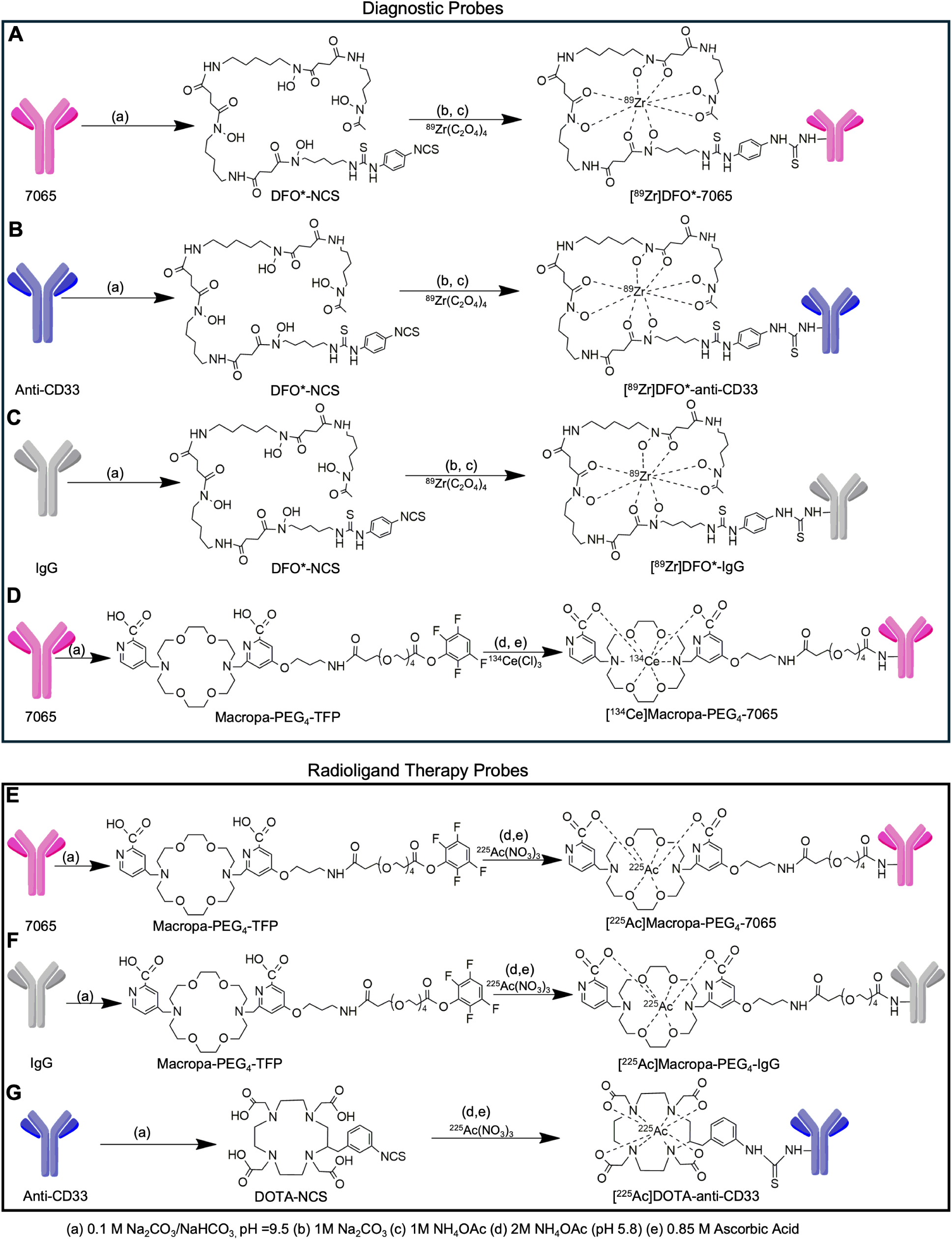
Schematic representation of radioimmunoconjugates. Three antibodies were employed, including 7065, which specifically targets the activated confirmation of aITGB2, anti CD33 (lintuzumab biosimilar), and non-targeting IgG, used as a control. These antibodies were conjugated with DFO*-NCS and ^89^Zr-oxalate to generate A) [^89^Zr]DFO*-7065, B) [^89^Zr]DFO*-anti-CD33 C) [^89^Zr]DFO*-7065, which were used for cell binding and PET imaging and biodistribution studies. 7065 was conjugated with Macropa-PEG_4_-TFP ester, and then with D)^134^Ce (for imaging) or E) ^225^Ac(NO_3_)_3_. (for therapy). F) The control IgG antibody was also conjugated with Macropa-PEG_4_-TFP and labelled with ^225^Ac(NO_3_)_3_ for therapy control studies. G) Anti-CD33 antibody (Lintuzumab) was conjugated DOTA-NCS and labeled with ^225^Ac(NO_3_)_3_.

For ^225^Ac therapy studies, we employed a recently designed bifunctional chelator, Macropa-PEG_4_-TFP (*26,27*). This bifunctional chelator employs the efficient Macropa chelator (*28*), together with a pegylated linker, which results in efficient radiolabeling, high tumor uptake, and rapid, renal clearance of metabolic fragments. This results in high tumor uptake and lower background tissue binding, with improved therapeutic outcomes compared to conventional DOTA-based labeling strategies (*27*). Utilizing optimized labeling protocols (*26*), Macropa-PEG_4_-7065 and Macropa-PEG_4_-IgG were obtained with antibody-to-chelator ratios of 1.61 and 0.43 (**Figure 2D-F, S7A-B**), respectively. The anti-CD33 antibody was conjugated to DOTA-NCS following a previously reported method, yielding an antibody-to-chelator ratio of 11.66 (**Figure 2G, S7C**). [^225^Ac]Macropa-PEG_4_-7065 and [^225^Ac]Macropa-PEG_4_-IgG were obtained with a radiochemical yield of 55 ± 1.76 % and 65 ± 0.06 % yield (n=4 syntheses each) with radiochemical purity exceeding 98% (**Figure S8A-D**) with a specific activity of 0.25 mCi/mg and 0.5 mCi/mg respectively. [^225^Ac]DOTA-anti-CD33 was obtained with a radiochemical yield of 40 ± 0.45 % yield (n = 4 syntheses) with a radiochemical purity of more than 98% (**Figure S8E-F**) with a specific activity of 0.166 mCi/mg. We developed a ^134^Ce labeled version of Macropa-PEG_4_-7065 to directly image this conjugate, achieving a 75 ± 0.32% radiochemical yield (n=3), with >95% purity and a specific activity of 5 mCi/mg (**Figure S8G-H**). SEC analysis for [^225^Ac]Macropa-PEG_4_-7065 (**Figure S9**) showed no aggregation of [^225^Ac]Macropa-PEG_4_-7065 after the labeling and purification process. Immunoreactivity was preserved for both [^225^Ac]Macropa-PEG_4_-7065 (78.6%) and [^225^Ac]DOTA-anti-CD33 (77.0%) with recombinant ITGB2 and CD33 proteins, respectively (**Figure S10A–B**). [^225^Ac]Macropa-PEG_4_-7065 remained >95% stable in saline and human serum for up to 7 days (**Figure S10C**). Overall, the radiosynthesis of the agents was robust, reproducible, with excellent purity, stability, specific activity, and retention of immunoreactivity.

### [^89^Zr]DFO*-7065 demonstrates high binding to AML cell lines, proportionate to the expression of aITGB2

We performed a cell-binding study to evaluate the binding of [^89^Zr]DFO*-7065 to different AML cell lines. Nomo-1 and HL-60 exhibited the highest cell-binding, followed by THP-1 and MV411, consistent with the higher expression of aITGB2 detected by flow cytometry. The binding was approximately 3-fold lower in the Nomo-1 ITGB2-KO cells as compared to the Nomo-1 WT cell line. Both Nomo-1 and Nomo-1 ITGB2-KO cells express Fc receptors. To block Fc receptor-mediated binding in the Nomo-1 KO cells, 10 fold excess of cold IgG was added, further reducing the binding of [^89^Zr]DFO*-7065 to the Nomo-1 ITGB2-KO cell line (**Figure 3A**). The dissociation constant (K_d_) of [^89^Zr]DFO*-7065 for all aITGB2-expressing cell lines (Nomo-1, HL-60, THP-1, MV411) was in the nanomolar range in between (2 nM – 60 nM) with a receptor density (B_max_) of (0.04 × 10^5^ – 0.25 × 10^5^) receptors/cell (**Figure 3B, S11**). The radiopharmaceutical displayed rapid internalization, with 52.9 ± 5.4% and 79.6 ± 3.6 % of total administered activity for Nomo-1 and HL-60, respectively (**Figure 3C**). These findings support the notion that aITGB2 is highly but variably expressed in AML cell lines and that [^89^Zr]DFO*-7065 binds selectively to the activated form of aITGB2 in these cells.

**Figure 3.**
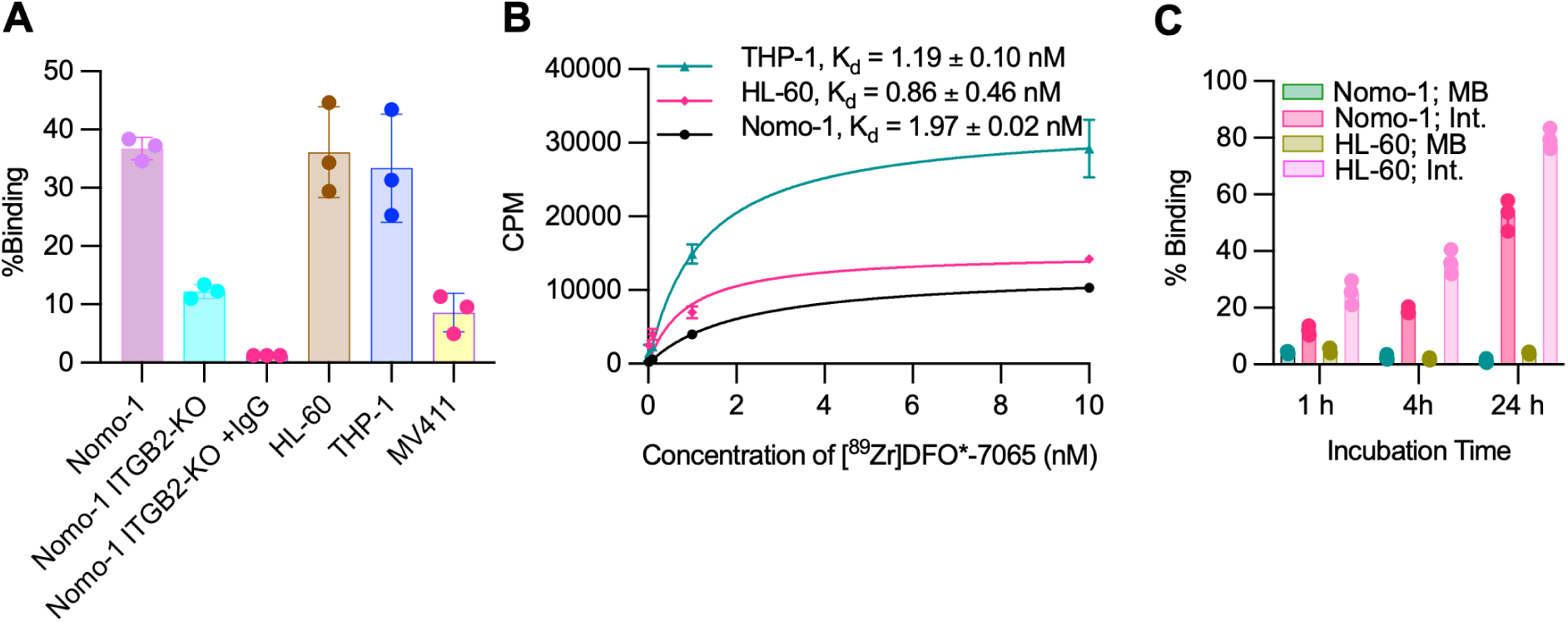
aITGB2 can be detected with [^89^Zr]DFO*-7065 in AML cell lines. A) Cell binding assay to measure the percentage cell-associated activity of [^89^Zr]DFO*-7065 using different AML cell lines B) K_d_ measurement of [^89^Zr]DFO*-7065 on Nomo-1, HL-60, and THP-1 cells C) Percentage internalized binding of [^89^Zr]DFO*-7065 with Nomo-1 and HL-60 cells.

### μPET/CT imaging and biodistribution analysis of [^89^Zr]DFO*-7065 in disseminated AML models reveal high aITGB2-targeted uptake

The imaging properties of [^89^Zr]DFO*-7065 were evaluated in disseminated AML models. AML cells were inoculated by tail vein, and bioluminescence and PET imaging were performed followed by biodistribution and imaging studies (**Figure 4A**). The μPET/CT imaging demonstrated high uptake of [^89^Zr]DFO*-7065 in the bone marrow of the Nomo-1 disseminated model, with spatial colocalization of bioluminescence (BLI) and PET signals (**Figure 4B**). To confirm the specificity of [^89^Zr]DFO*-7065 targeting aITGB2, we performed three control experiments including imaging with [^89^Zr]DFO*-7065 with blocking using a 25-fold excess of unlabeled 7065 (**Figure 4C**), imaging with a non-specific [^89^Zr]-labeled antibody ([^89^Zr]DFO*-IgG; **Figure 4D**), and imaging with [^89^Zr]DFO*-7065 in the Nomo-1 ITGB2-KO model (**Figure 4E**). All three studies showed significantly reduced uptake in tumor lesions, with no overlap between the BLI and PET/CT signals, confirming the specificity of [^89^Zr]DFO*-7065 for aITGB2. Region of interest (ROI) analysis confirmed higher uptake of [^89^Zr]DFO*-7065 in femur and excretion by liver (**Figure S 12A-D**). *Ex vivo* BLI and PET images demonstrated high concordance of signal for Nomo-1, but not the control studies (**Figure 4F, S12E-H**). *Ex vivo* biodistribution studies showed significantly higher uptake of [^89^Zr]DFO*-7065 in the bone marrow as compared to negative controls (**Figure 4G, S12I**). The Tumor/Blood (**Figure 4H**) and Tumor/Muscle (**Figure 4I**) ratios were significantly higher in the Nomo-1 model compared to the control models. These results confirm that [^89^Zr]DFO*-7065 has high binding in the Nomo-1 disseminated model, with specific target mediated binding to aITGB2.

**Figure 4.**
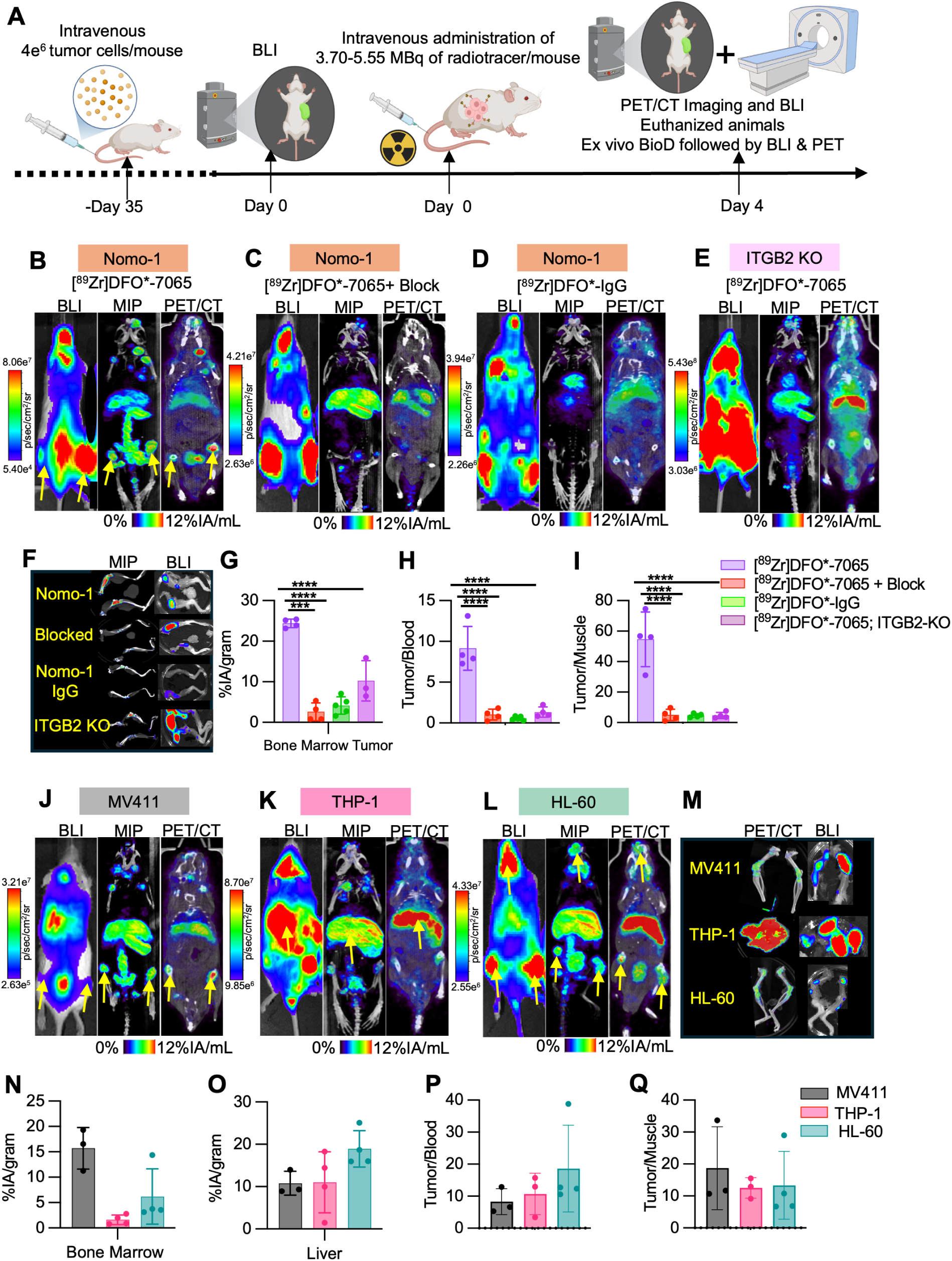
[^89^Zr]DFO*-7065 effectively detects aITGB2 positive AML disseminated lesions with high target to background PET imaging. A) Schematic of in vivo and *ex vivo* imaging and biodistribution studies in disseminated AML models. B) BLI, MIP, and μPET/CT fusion images of in the Nomo-1 disseminated with [^89^Zr]DFO*-7065, with high tumoral uptake, reduced with C) co-administration of a 25-fold excess of 7065, or D) when non-targeting [^89^Zr]DFO*-IgG was administered. E) Low uptake of [^89^Zr]DFO*-7065 was observed in the Nomo-1 ITGB2-KO model. F) *Ex vivo* BLI and μPET/CT fusion images of femurs harvested from Nomo-1-Luc and Nomo-1 ITGB2-KO-Luc models. G) *Ex vivo* biodistribution indicates significantly higher tumor uptake as well as H) increased tumor/blood ratio and I) tumor/muscle ratios in the [^89^Zr]DFO*-7065 Nomo-1 group compared to controls. J) BLI and μPET/CT fusion images of [^89^Zr]DFO*-7065 in MV411-Luc K) in THP-1-Luc, and in L) HL-60-Luc, all with high tumoral uptake. M) *Ex vivo* BLI and MIP images of [^89^Zr]DFO*-7065 in MV411-Luc, HL-60-Luc, THP-1-Luc model in femur (for MV411 and HL-60) and liver (for THP-1), with spatial colocalization of tumoral BLI and radiopharmaceutical uptake. N) *Ex vivo* biodistribution of [^89^Zr]DFO*-7065 in bone marrow and O) liver, revealing high uptake in the tumor-containing organs. P) Tumor/blood and Q) tumor/muscle ratio of [^89^Zr]DFO*-7065 in all groups at 4 days post injection.

We extended this evaluation to the MV411, THP-1, and HL-60 AML disseminated models. BLI showed variable anatomic leukemic cell accumulation: Nomo-1 and HL-60 localized primarily to bone marrow, THP-1 to liver, and MV411 to both liver and bone marrow (**Figure 4J–L**). In all cases, [^89^Zr]DFO*-7065 localized concordantly with the tumor-associated BLI signal. *Ex vivo* μPET/CT and BLI analyses of bone marrow confirmed signal colocalization across models (**Figure 4M**). *Ex vivo* analyses also showed high tumor cell accumulation in spleen and lungs in all three models, with concordant [^89^Zr]DFO*-7065 uptake (**Figure S12J–L**). Biodistribution studies showed similar findings (**Figure 4N, S12M**). Tumor/blood (**Figure 4P**) and tumor/muscle (**Figure 4Q**) ratios were high across all models. ROI analysis performed at day 4 demonstrate high uptake of [^89^Zr]DFO*-7065 in femur and liver concordance with tumor signal in BLI and minimal uptake in the kidney and blood pool (**S12N-Q**). Overall, these findings revealed a strong tumor signal with low background, and supporting the further evaluation of therapeutic radiopharmaceuticals.

To compare against the current standard of care imaging methods, μPET/CT imaging and biodistribution studies were performed using [^18^F]-FDG in Nomo-1 and THP-1 disseminated models. μPET/CT imaging revealed some [^18^F]-FDG uptake in tumors (**Figure S13A, S13B**), which corresponded to areas identified by BLI. However, the uptake in tumor sites, particularly in the femur and liver, was significantly lower than in other background organs including the heart, kidneys, and brain. ROIs drawn for the femur and liver demonstrated reduced [^18^F]-FDG accumulation (**Figure S13C, S13D**), while kidneys and heart exhibited higher uptake (**Figure S13E, S13F**) consistent with their roles in [^18^F]-FDG metabolism and excretion. *Ex vivo* biodistribution studies corroborated these findings, showing a lower %IA/g of [^18^F]-FDG in bone marrow tumors compared to other organs (**Figure S13G**), and lower tumor/blood and tumor/muscle ratio (**Figure S13H, S13I**) compared to [^89^Zr]DFO*-7065. Overall, [^89^Zr]DFO*-7065 demonstrated superior imaging properties, with higher tumor uptake and tumor to background ratios, when compared to standard of care [^18^F]-FDG.

### [^225^Ac]Macropa-PEG_4_-7065 demonstrates aITGB2 dependent cell killing *in vitro* and favorable tumoral and whole animal biodistribution *in vivo*

After evaluating the imaging capability of [^89^Zr]DFO*-7065, we analyzed the therapeutic efficacy of [^225^Ac]Macropa-PEG_4_-7065 *in vitro*. In clonogenic survival assays, dose dependent reduction in colonies was observed after treatment with [^225^Ac]Macropa-PEG_4_-7065 in Nomo-1 cells (IC_50_ of 90.05 ± 0.03 pCi/ml) with attenuated efficacy in Nomo-1 ITGB2-KO cells (3.08 ± 0.90 nCi/mL) (**Figure 5A**). In the highly aITGB2 expressing HL-60 cell line, the IC_50_ was 80.3 ± 0.03 pCi/mL and in the moderately expressing MV411 cell line the IC_50_ was 10.61 ± 4.74 nCi/mL. These findings recapitulate the trends seen in flow cytometry (**Figure 1A**) and cell binding (**Figure 3A**) studies, linking the expression levels of aITGB2 to therapeutic efficacy.

**Figure 5.**
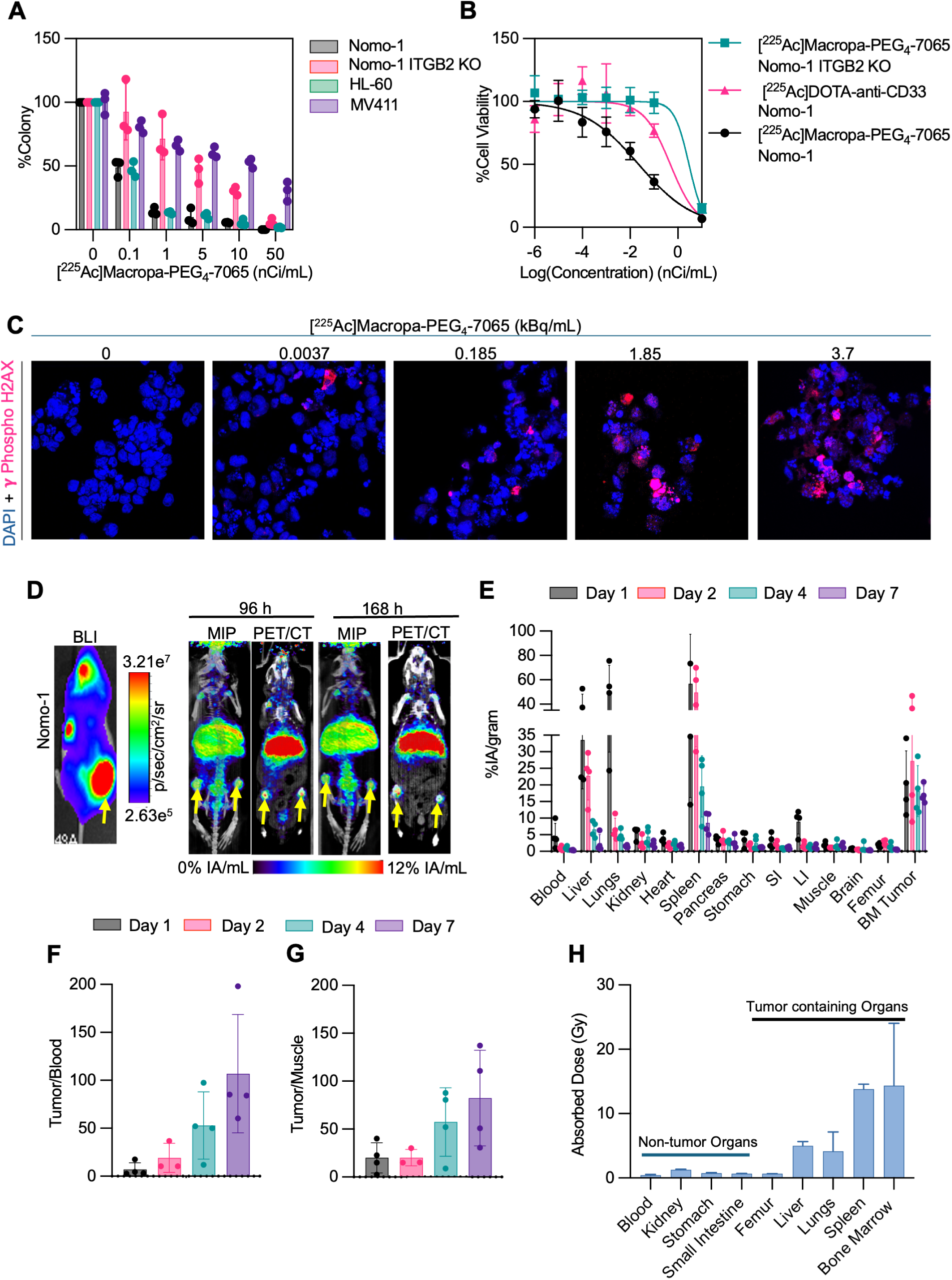
[^225^Ac]Macropa-PEG_4_-7065 demonstrates high therapeutic efficacy and [^134^Ce/^225^Ac]Macropa-PEG_4_-7065 demonstrates favorable tumoral and whole animal biodistribution. A) Colony formation assay using [^225^Ac]Macropa-PEG_4_-7065 in Nomo-1 and Nomo-1 ITGB2-KO, HL-60 and MV411 cell lines. B) Cell killing/viability assay with [^225^Ac]Macropa-PEG_4_-7065 in Nomo-1 and Nomo-1 ITGB2-KO cell line and with [^225^Ac]DOTA-anti-CD33 in Nomo-1 cell line C) γ-phospho H2AX staining for DNA damage analysis following [^225^Ac]Macropa-PEG_4_-7065 treatment revealed dose-dependent increase in the H2AX foci. D) BLI, MIP and μPET/CT fusion images of [^134^Ce]Macropa-PEG_4_-7065 in Nomo-1-Luc bearing mouse at 96 h and 168 h post injection. E) *Ex vivo* biodistribution study in Nomo-1-Luc model indicating accumulation of [^225^Ac]Macropa-PEG_4_-7065 in % IA/gram. F) Tumor/Blood and H) Tumor/Muscle ratio at different time points of [^225^Ac]Macropa-PEG_4_-7065, indicating higher accumulation and retention of [^225^Ac]Macropa-PEG_4_-7065 as compared to blood pool and background organs. G) Dosimetry analysis of biodistribution data demonstrates high absorbed dose in tumor bearing organs, but not healthy tissues.

For comparison of the therapeutic efficacy of [^225^Ac]Macropa-PEG_4_-7065 and to serve as a positive control, we utilized a previously reported radioimmunoconjugate [^225^Ac]DOTA-anti-CD33 (biosimilar for Lintuzumab) which is now in clinical trials for the treatment of AML (*29,30*). The K_d_ value for anti-CD33 antibody with CD33 recombinant protein was measured as 1.16 ± 0.005 nM by biolayer interferometry (**Figure S14**), similar to the value reported in previous studies (*31*). We performed a cytotoxicity assay to compare the therapeutic efficacy of [^225^Ac]Macropa-PEG_4_-7065 with [^225^Ac]DOTA-anti-CD33. [^225^Ac]Macropa-PEG_4_-7065 showed greater efficacy, with a significantly lower IC_50_ (0.022 ± 0.02 nCi/mL) compared to [^225^Ac]DOTA-anti-CD33 with a IC_50_ value of 0.40 ± 0.25 nCi/mL (*p* < 0.05) (**Figure 5B**). In contrast, the IC_50_ value for [^225^Ac]Macropa-PEG_4_-7065 in Nomo-1 ITGB2-KO cells was markedly higher (4.42 ± 2.99 nCi/mL), demonstrating that aITGB2 is necessary for a strong therapeutic effect. In addition, a dose-dependent increase of phospho-γ-H2AX foci, indicating DNA damage, was observed after [^225^Ac]Macropa-PEG_4_-7065 treatment (**Figure 5C, S15**). These data reveal that [^225^Ac]Macropa-PEG_4_-7065 is effective for killing AML cells *in vitro* in a manner that requires aITGB2, with greater therapeutic efficacy compared to [^225^Ac]DOTA-antiCD33.

After observing promising therapeutic efficacy *in vitro* in AML cell lines, we evaluated the biodistribution and dosimetry of [^225^Ac/^134^Ce]Macropa-PEG_4_-7065 *in vivo* in an AML disseminated model. In this case, [^134^Ce]Macropa-PEG_4_-7065 was used as a surrogate of [^225^Ac]Macropa-PEG_4_-7065 to monitor pharmacokinetics and biodistribution analysis, as has been previously reported for other radiopharmaceuticals (*26*). As expected, BLI and μPET/CT imaging demonstrated high, and gradually increasing [^134^Ce]Macropa-PEG_4_-7065 in the bone marrow over 7 days, with spatial colocalization of bioluminescence and PET signals (**Figure 5D**). *Ex vivo* μPET/CT and BLI showed the matching areas of uptake of [^134^Ce]Macropa-PEG_4_-7065 in femur, liver, and spleen (**Figure S16A)**. In addition, we performed a multi-time point *ex vivo* biodistribution for [^225^Ac]Macropa-PEG_4_-7065 in the Nomo-1 disseminated model, which revealed high and sustained uptake in bone marrow tumor, with favorable tumor to blood and tumor to muscle ratios (**Figure 5E-G**). The *ex vivo* Biodistribution studies performed 7 days post injection of [^134^Ce]Macropa-PEG_4_-7065 and [^225^Ac]Macropa-PEG_4_-7065 showed that the uptake of both the radioimmunoconjugates in bone marrow and other organs was similar except in liver and spleen (**Figure S16B**). Dosimetry analysis of [^225^Ac]Macropa-PEG_4_-7065 revealed that tumor-infiltrated organs including bone marrow, liver, and spleen and lungs tumor showed high absorbed dose, compared to non-tumor bearing tissues (**Figure 5H**). These results suggest that [^225^Ac]Macropa-PEG_4_-7065 exhibits high specificity and potent cytotoxicity against AML cell lines, and a favorable biodistribution, suggesting feasibility for therapeutic studies.

### Despite Murine aITGB2 Cross-Reactivity, [^225^Ac]Macropa-PEG_4_-7065 Does Not Exacerbate Bone Marrow Suppression Beyond Non-Specific ^225^Ac Effects

After validating promising therapeutic efficacy *in vitro*, we evaluated the toxicity profile and maximum tolerated activity of [^225^Ac]Macropa-PEG_4_-7065. Initial acute toxicity studies of 9.25 kBq of [^225^Ac]Macropa-PEG_4_-7065 were carried out at day 7 and day 28 post administration in healthy black C57BJ/6J mice and compared with vehicle group injected with saline, 9.25 kBq of [^225^Ac]Macropa-PEG_4_-IgG and 9.25 kBq of [^225^Ac]DOTA-anti-CD33 injected cohorts (**Figure S17A**). It is important to note that we previously found our 7065 clone that recognizes human aITGB2 in a conformation-selective manner cross-reacts with murine aITGB2; however, the lintuzumab anti-CD33 antibody is not thought to be murine cross-reactive (*32*). At day 7 and 28, white blood cells, lymphocytes, monocytes, blood platelets and basophils showed significant decrease in cell count in [^225^Ac]Macropa-PEG_4_-7065 and [^225^Ac]Macropa-PEG_4_-IgG injected cohort as well as in [^225^Ac]DOTA-anti-CD33 injected group, as compared to the vehicle injected group. Other blood parameters including hemoglobin and platelet counts were similar between groups (**Figure S17(a-g) and (h-v)**). Histopathology of dose-limiting organs (kidney, liver, spleen, lungs, bone marrow, and heart) showed that all organs were normal in all ^225^Ac-radiopharmaceuticals injected cohorts, except for the kidney where there was minimal renal focal tubular damage. The tubular damage in the kidneys for all cohorts was similar compared to the vehicle group (**Figure S18**).

A chronic toxicity study of [^225^Ac]Macropa-PEG_4_-7065 was performed in NRG mice with an endpoint at day 100 post administration, using both single as well as fractionated administration regimens (**Figure S19A**). No significant body weight loss or survival difference (**Figure S19B, S19C**), or hematologic or metabolic panel abnormalities (**Figure S19D-O)** were observed in any cohort. Histopathology of dose-limiting organs showed that in the case of single and fractionated doses of 4.62 kBq of [^225^Ac]Macropa-PEG_4_-7065 injected groups, all organs were normal, except in the case of two of 5 mice, where minimal renal focal tubular damage was observed in the kidney (**Figure S20**). Similarly, in case of single and fractionated dose of 9.25 kBq of [^225^Ac]Macropa-PEG_4_-7065 injected groups, minimal damage to kidney was observed with renal injury in one of 5 mice (single dose) and all of the 5 mice (fractionated dose), presumably due to redistribution of daughter ^213^Bi into the kidney, as previously reported (*33*). Taken together, these findings suggest that [^225^Ac]Macropa-PEG_4_-7065 demonstrates the expected toxicity profile for an ^225^Ac-IgG, with transient hematologic abnormalities and delayed nephrotoxicity, with no additional marrow toxicity due to “on-target, off tumor” hematopoietic cell targeting.

### PET/CT imaging in hCD34⁺ humanized immune system mice reveals higher binding of [^89^Zr]DFO*-CD33 to the human immune system compared to [^89^Zr]DFO*-7065

In order to further verify the specificity of the 7065 antibody and analyze the toxic effect of 7065 and anti-CD33 radioimmunoconjugates to bone marrow and human immune cells, we performed μPET/CT imaging in hCD34^+^ humanized immune system (NOG EXL) mice injected with [^89^Zr]DFO*-anti-CD33 and [^89^Zr]DFO*-7065. We observed high background binding of [^89^Zr]DFO*-anti-CD33 to human CD45+ immune cells (**Figure S21A**) as compared to [^89^Zr]DFO*-7065 (**Figure S21B**) at day 7 post-injection. ROI analysis drawn on femur also confirmed higher binding of [^89^Zr]DFO*-anti-CD33 to human immune cells as compared to [^89^Zr]DFO*-7065 at day 7 (p < 0.045) (**Figure S21C**). These findings confirm lower binding to normal human bone marrow and human immune cells for [^89^Zr]DFO*-7065 compared to [^89^Zr]DFO*-anti-CD33, suggesting potential for reduced bone marrow toxicity and improved therapeutic index in future clinical studies.

### [^225^Ac]Macropa-PEG_4_-7065 demonstrates high therapeutic efficacy and prolonged survival in Nomo-1 disseminated AML model

A treatment study was designed to evaluate the therapeutic efficacy of [^225^Ac]Macropa-PEG_4_-7065 (**Figure 6A**), with groups including vehicle control, non-targeting [^225^Ac]Macropa-PEG_4_-IgG, the previously evaluated leukemia RIT [^225^Ac]DOTA-anti-CD33, and [^225^Ac]Macropa-PEG_4_-7065. An additional group received three fractionated doses of [^225^Ac]Macropa-PEG_4_-7065. In the vehicle group as well as the [^225^Ac]Macropa-PEG_4_-IgG-injected group, a continuous increase in tumor burden was observed on BLI, with all mice reaching the endpoint at day 43 and day 50, respectively (**Figure 6B**). In the [^225^Ac]DOTA-anti-CD33 cohort, there was an initial decrease in tumor burden during the first week, followed by a continuous increase, with all mice reaching the endpoint at day 43. In contrast, the single-dose and fractionated-dose groups of [^225^Ac]Macropa-PEG_4_-7065 demonstrated a significant decrease in tumor burden. No weight loss was observed in any of the cohort, except one mouse injected with a total three fractionated doses of 9.25 kBq (a non-statistically significant trend with approximate ≤ 5% body weight loss) (**Figure 6C**). In the single-dose [^225^Ac]Macropa-PEG_4_-7065 injected group, two mice relapsed, showing high tumor burden, and were euthanized at days 57 and 71, respectively. However, the six out of eight remaining animals in this group survived until the end of the study, which was terminated on day 148. The median survival time for the vehicle group was 43 days, 50 days for [^225^Ac]Macropa-PEG_4_-IgG, and 43 days for the [^225^Ac]DOTA-anti-CD33 group. In contrast, for both the single-dose and fractionated-dose [^225^Ac]Macropa-PEG_4_-7065 injected groups, the majority of animals survived in this long-term study; therefore, the median survival was undefined and was significantly higher than negative and positive controls (**Figure 6D**).

**Figure 6.**
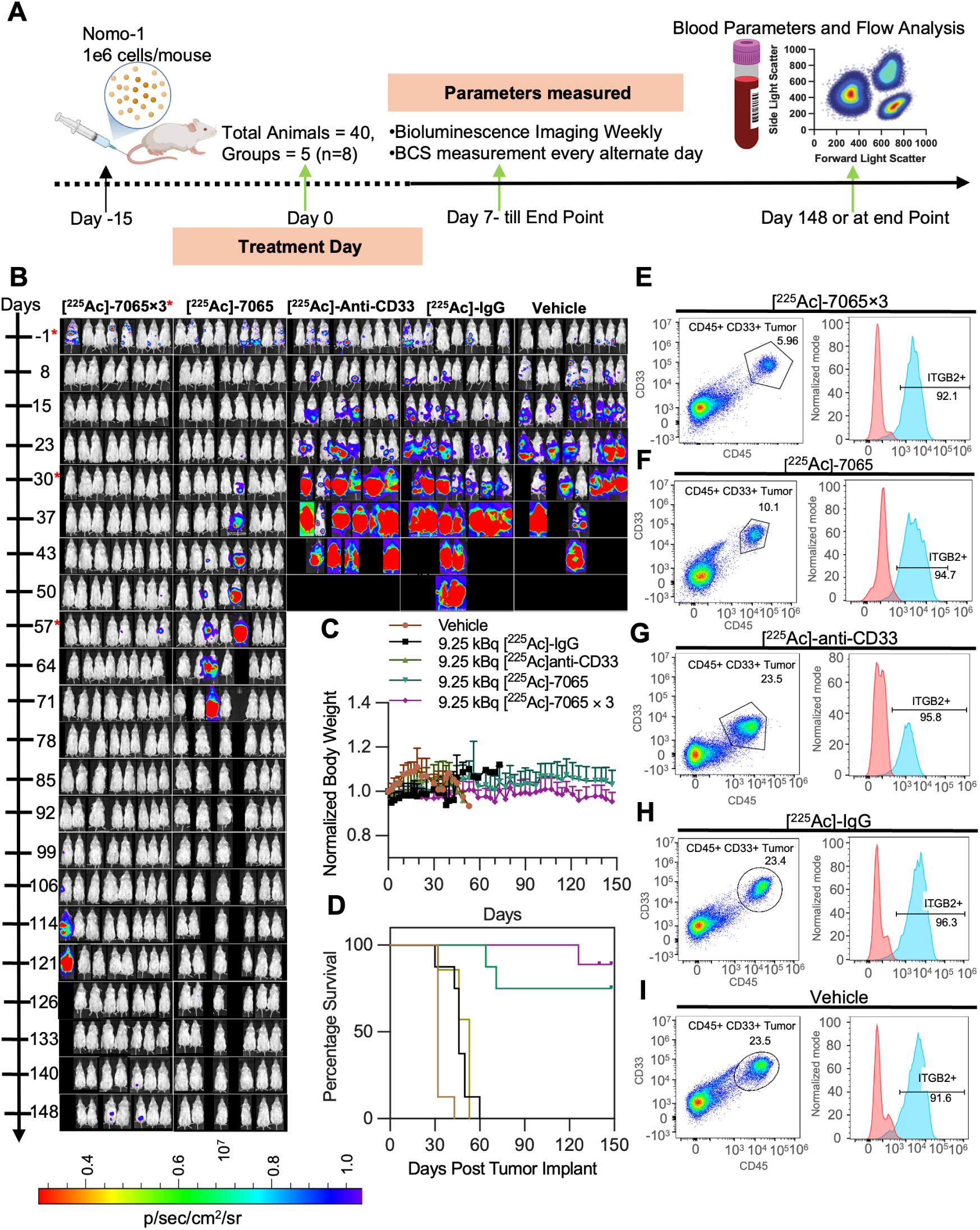
[^225^Ac]Macropa-PEG4-7065 based radiopharmaceutical therapy for effective treatment of AML. A) Schematic showing the workflow for the therapy study. B) Serial BLI imaging indicates reduced tumor burden in the [^225^Ac]Macropa-PEG_4_-7065 single and fractionated administration groups (dosed on day 0, day 30, and day 57) as compared to saline, [^225^Ac]Macropa-PEG_4_-IgG and [^225^Ac]DOTA-anti-CD33 cohorts. C) Body weight measurements D) Kaplan Meier curve demonstrates improvement in overall survival in the [^225^Ac]Macropa-PEG_4_-7065 groups. FACS analysis indicating CD45+ CD33+ tumor population and aITGB2 expression for a representative mouse injected with E) fractionated administration or F) single administration of [^225^Ac]Macropa-PEG_4_-7065 G) [^225^Ac]DOTA-anti-CD33 H) [^225^Ac]Macropa-PEG_4_-IgG or I) saline + IgG.

We monitored aITGB2 expression in relapsed tumors to investigate potential resistance to treatment. CD33 and CD45-positive cells were isolated from the bone marrow and spleen when animals approached the endpoint, and aITGB2 expression in the CD33/CD45-positive tumor population was analyzed using flow cytometry (**Figure S22**). Interestingly, aITGB2 expression remained high (greater than 95%) in the CD33/CD45-positive tumor population in bone marrow, including single-or fractionated-dose [^225^Ac]Macropa-PEG_4_-7065 groups (**Figure 6E-F, S22A-B**), [^225^Ac]DOTA-anti-CD33 (**Figure 6G, S22C-D**), [^225^Ac]Macropa-PEG_4_-IgG (**Figure 6H, S22E-F**), saline + IgG (**Figure 6I, S22G-H**) injected groups. Similarly, aITGB2 expression remained high in the CD33/CD45-positive tumor population in spleen across all the cohorts (**Figure S23A-L**). These results illustrate that [^225^Ac]Macropa-PEG_4_-7065 demonstrates promising therapeutic efficacy compared to control groups as well as [^225^Ac]DOTA-anti-CD33. Furthermore, persistent antigen expression suggests the potential for additional administrations to further improve therapeutic efficacy.

### The proteome of relapsed tumors is remodeled in response to [^225^Ac]Macropa-PEG_4_-7065

For the mice found to have relapsed tumors in our Nomo-1 *in vivo* study (**Figure 6**), we performed comprehensive proteomic analysis to understand the molecular mechanism of tumor remodeling after aITGB2 [^225^Ac]Macropa-PEG_4_-7065 treatment. Comparison between [^225^Ac]Macropa-PEG_4_-7065 based versus [^225^Ac]Macropa-PEG_4_-IgG isotype control groups revealed extensive molecular remodeling in response to therapy, with 558 proteins significantly upregulated and 373 significantly downregulated (*p*-value <0.05, |Log_2_FC| > 1.5) (**Figure 7A, S24**), and differential protein identifications in [^225^Ac]Macropa-PEG_4_-7065 treated mice vs. IgG or saline control. 1,515 proteins were uniquely detected in the [^225^Ac]Macropa-PEG_4_-7065 group (**Figure 7B**), highlighting therapy-induced proteomic changes. Hierarchical clustering showed a coordinated upregulation of specific protein groups in the [^225^Ac]Macropa-PEG_4_-7065 treated samples (**Figure 7C**), where pathway enrichment analysis (**Figure 7D**) revealed strong activation of the proteasome machinery, TCA cycle, carbon metabolism, and amino acid metabolism, likely reflecting a cellular adaptation to therapy-induced stress. Furthermore, pathways associated with immune recognition including cell adhesion, phagosome, antigen processing and presentation, and various N-glycan biosynthesis processes were significantly downregulated (**Figure 7E**), potentially indicating altered interactions with the tumor microenvironment as a mechanism of resistance to [^225^Ac]Macropa-PEG_4_-7065 RIT. Notably, N-glycan biosynthesis and antigen processing and presentation were among the most suppressed (FDR < 10⁻⁴), along with phagosome formation, cell adhesion molecules, mucin-type O-glycan biosynthesis, and glycosaminoglycan biosynthesis.

**Figure 7.**
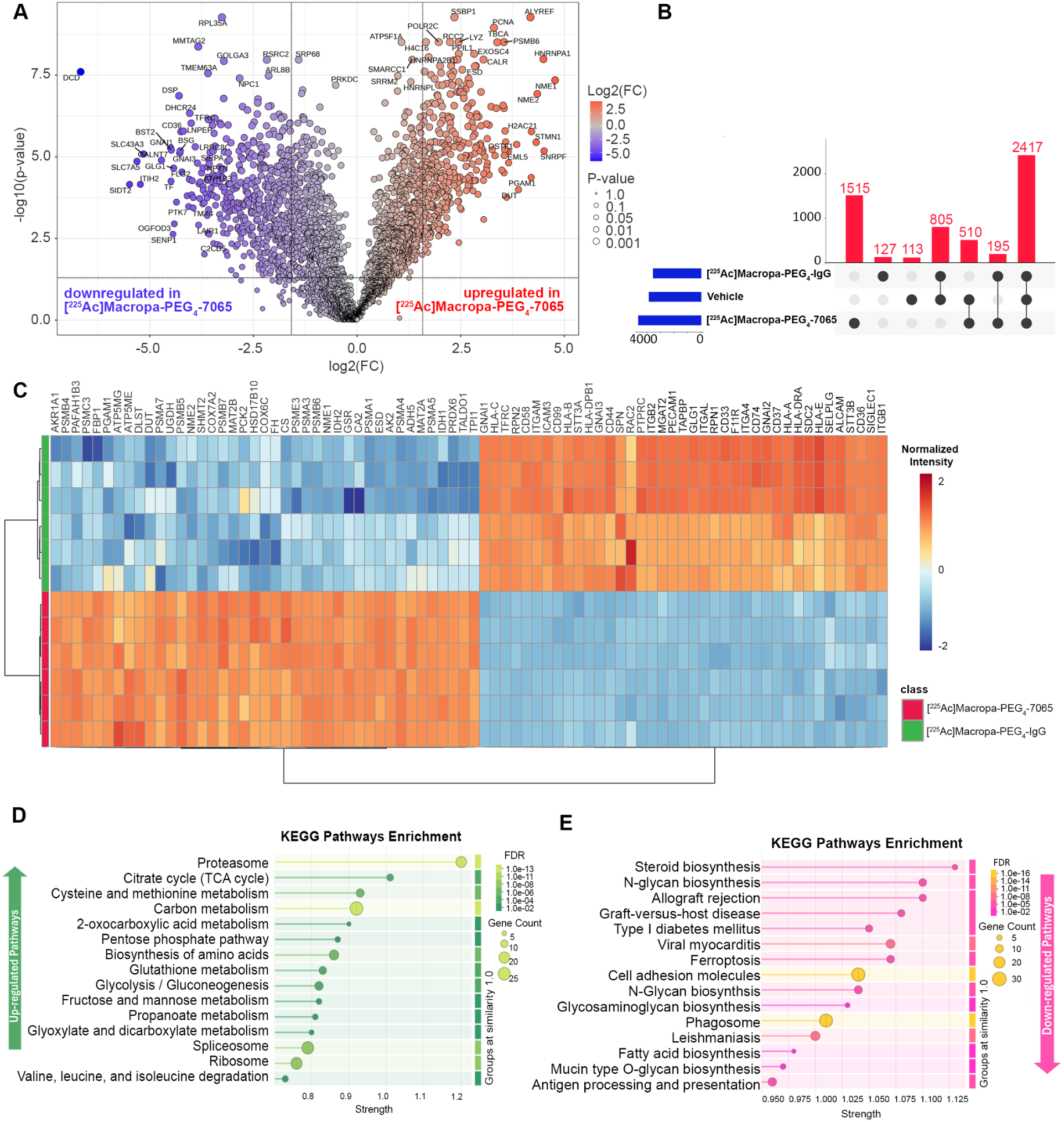
Global proteomic analysis following radioimmunotherapy performed in relapsed mice. A) Volcano plot showing differential protein expression between [^225^Ac]Macropa-PEG_4_-7065–treated AML cells and [^225^Ac]Macropa-PEG_4_-IgG. B) The UpSet plot shows protein identifications across experimental groups. C) Heatmap of the top differentially expressed proteins shows clear segregation between treatment and control groups, with distinct clusters reflecting coordinated shifts in metabolic, translational, and immune pathways. D) KEGG enrichment analysis of upregulated proteins revealed strong activation of key metabolic and proteostatic programs. E) In contrast, downregulated proteins were significantly enriched for pathways associated with immune recognition and glycan processing.

### [^225^Ac]Macropa-PEG_4_-7065 is an effective treatment in a patient derived xenograft model of AML

We next evaluated the therapeutic efficacy of [^225^Ac]Macropa-PEG_4_-7065 in a disseminated PDX model of AML (obtained from ProXe biobank(*21*)) implanted in NRG-SGM3 mice (n=5/group). Following preconditioning with busulfan and PDX tumor inoculation, treatment was initiated on day 10 with treatment arms including vehicle, [^225^Ac]DOTA-anti-CD33, and [^225^Ac]Macropa-PEG_4_-7065 group (**Figure 8A**). Efficacy was assessed by flow cytometry of human CD45⁺ (hCD45^+^) cells in blood, bone marrow, and spleen, along with body weight and condition scoring. At day 15, minimal or no hCD45^+^ population was observed in all cohorts by flow analysis (**Figure S25A-C**). By day 30, high tumor burden was observed in the vehicle (saline) group, with ∼90% hCD45⁺ blasts detected by flow cytometry (**Figure 8B, S26A**). Significantly lower burden was observed in both the [^225^Ac]DOTA-anti-CD33 cohort and [^225^Ac]Macropa-PEG_4_-7065 cohort, with the latter group showing less than 2% hCD45⁺ blasts in all mice (**Figure 8C-D, S26B-C**). By day 41, all mice treated with [²²⁵Ac]DOTA-anti-CD33 showed high tumor burden and reached endpoint (**Figure 8E, S27A**). In contrast, [²²⁵Ac]Macropa-PEG_4_-7065–treated mice had lower tumor burden, with only two mice showing >10% hCD45⁺ blasts (**Figure S27B**). However, all relapsed injected with [²²⁵Ac]Macropa-PEG_4_-7065 by day 67, with three reaching end-point on day 52 and the rest by day 67 (**Figure S27C, S27D**). No body weight loss was seen at early time points following [^225^Ac]Macropa-PEG_4_-7065 treatment, while loss at later time points is likely due to tumor relapse (**Figure 8F**). Average overall survival was 30 days, 43 days, and 67 days in the saline, [^225^Ac]DOTA-anti-CD33, and [^225^Ac]Macropa-PEG_4_-7065 cohorts respectively (**Figure 8G**). Upon reaching the endpoint, bone marrow and spleen were harvested to assess hCD45⁺ cell levels. In all groups, high hCD45⁺ tumor population were observed in bone marrow and spleen (**Figure S28A-F)**. These results further confirmed in a patient-relevant model the promising therapeutic potential of [^225^Ac]Macropa-PEG_4_-7065.

**Figure 8.**
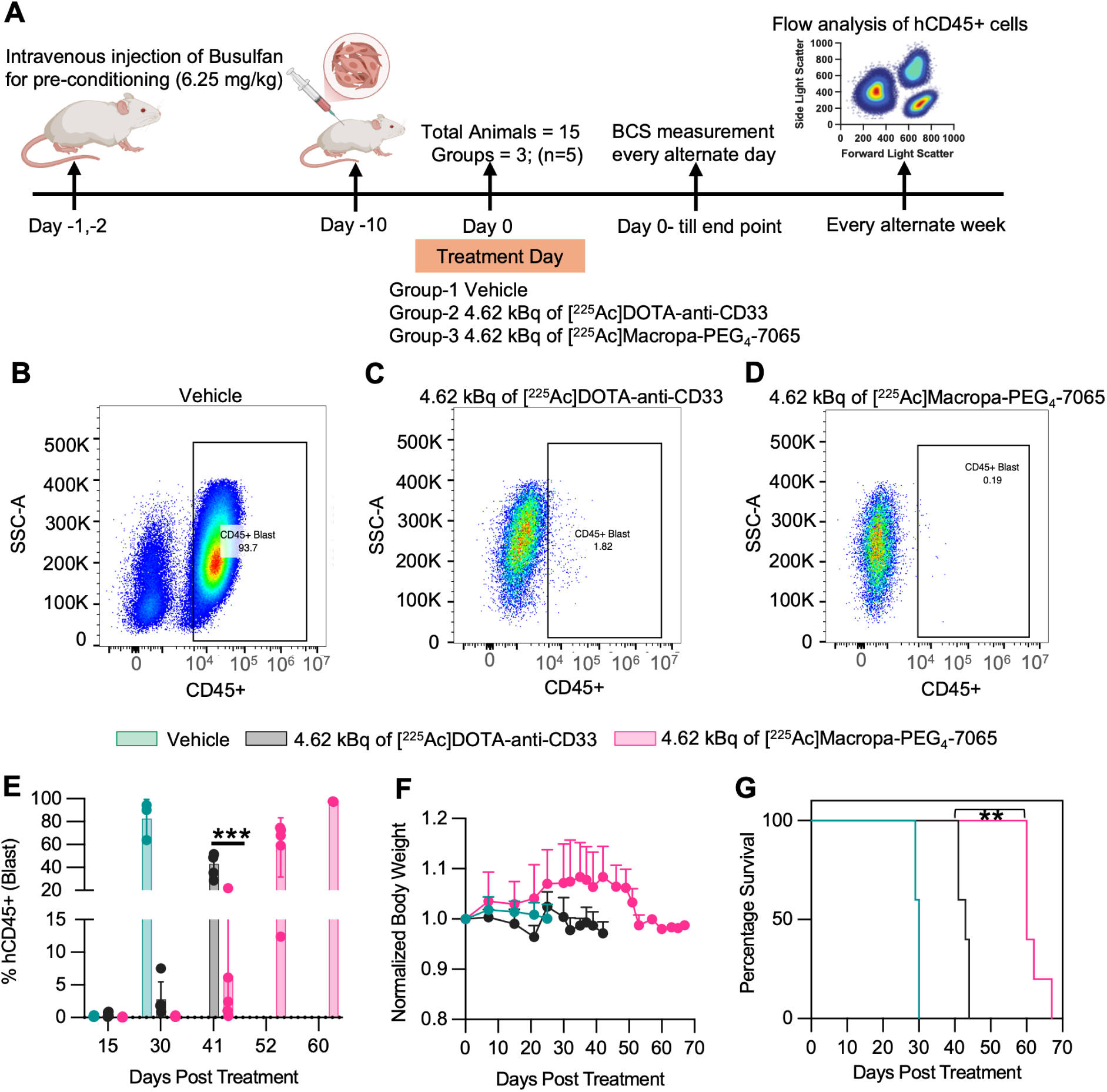
[^225^Ac]Macropa-PEG_4_-7065 based radiopharmaceutical therapy for the effective treatment of Patient Derived Xenografts. A) Schematic showing the workflow for the therapy study. Flow analysis of peripheral blood at day 30 for one representative mouse in B) saline C) [^225^Ac]DOTA-anti-CD33 and D) 4.62 kBq of [^225^Ac]Macropa-PEG_4_-7065 groups indicating CD45+ tumor population. E) Fraction of CD45+ blasts in peripheral blood in all the groups at day 15, day 30, day 41, day 52 and day 60 after treatment. F) Body weight measurement. G) Kaplan Meier curve demonstrates statistically significant improvement in overall survival in the [^225^Ac]Macropa-PEG_4_-7065 group.

## Discussion

AML is a heterogeneous malignancy that primarily affects individuals over 65 and remains difficult to cure despite the development of targeted therapies (*2,34*). Surface antigen-targeted treatments, such as antibody-based (*35*) and CAR T-cell therapies (*36*), show promise but are hindered by a lack of AML-specific targets, myelosuppression, and antigen escape. Through surface proteomics, we identified the activated conformation of integrin-β2 as a promising AML-specific target, and a conformation-selective antibody (7065) was developed and engineered into CAR T-cells, demonstrating strong efficacy in preclinical models (*21*). However, the challenges of CAR T-cell therapy in AML necessitate alternative strategies (*37*). Radioligand therapy (RLT) offers such an option by using antibodies to deliver radiation directly to leukemic cells, inducing targeted cytotoxicity with minimal damage to healthy tissues (*38*). Unlike conventional therapies, RLT is not reliant on immune activation and is less susceptible to antigen loss effective even in antigen-negative tumor cells due to the “crossfire effect,” making it viable option for patients with relapsed or refractory AML by targeting dispersed cells deep within the bone marrow.

With previously described targets and modalities, RIT in AML has shown limited clinical response with no FDA-approved products to date (*18*). Early studies using beta-particle emitters such as Iodine-131 and Yttrium-90 conjugated to anti-CD33 antibodies (M195 and huM195) demonstrated some reduction in leukemic burden, but their therapeutic impact was modest, especially in patients with high tumor loads (*39–41*). The long path length and lower linear energy transfer (LET) of beta emitters caused nonspecific damage to bystander cells, leading to prolonged myelosuppression and often requiring hematopoietic stem cell transplantation. These challenges have driven the exploration of alpha-particle emitters like Bismuth-213 and Actinium-225 conjugated to huM195 (Lintuzumab). Alpha emitters deposit higher energy over much shorter distances, allowing for potent and precise cell killing with less off-target toxicity, reduced myelosuppression, and the potential for use outside of transplant settings (*42–44*).

In this study, we have presented the systemic evaluation of aITGB2-targeted PET/CT imaging with [^89^Zr]DFO*-7065 and radioligand therapy with [^225^Ac]Macropa-PEG_4_-7065 in disseminated models of AML. This is the first study that explored the potential of aITGB2 as a radioimmunotherapy target for the diagnosis and treatment of AML. Flow cytometry indicated that the majority of tumors have high expression of aITGB2, and, importantly, antigen expression is independent of current molecular markers of AML classification. PET imaging revealed high specific uptake of [^89^Zr]DFO*-7065 in all disseminated AML models (Nomo-1, MV411, THP-1, HL-60). Relevant control studies including aITGB2 knock out, non-specific IgG imaging, and blocking demonstrate that the uptake of [^89^Zr]DFO*-7065 is specific to aITGB2 expression and not dominated by non-specific accumulation. Moreover, [^89^Zr]DFO*-7065 demonstrated higher tumor uptake and improved tumor to background ratios compared to the standard of care [^18^F]-FDG. These results demonstrate that [^89^Zr]DFO*-7065 is an effective agent that specifically binds aITGB2, and can be employed for detecting AML in preclinical models. While molecular imaging is less commonly employed in the management of leukemia in the clinic in comparison with solid tumors or lymphoma, this agent could find future use for dosimetry analysis to advance a companion radiopharmaceutical therapy or as a biomarker for selection of patients for subsequent aITGB2 targeted therapy. Overall, the collective data support the need for further therapeutic evaluation in preclinical studies and in the clinic.

[^225^Ac]-Lintuzumab, an alpha-emitting radioimmunotherapy agent directed against CD33-positive AML cells, has been evaluated in both preclinical and clinical studies (*11–12, 29–30*). Preclinical research demonstrated its potent cytotoxic effects due to alpha-particle–induced DNA damage, making it an effective option for eliminating leukemia cells, even in cases resistant to conventional therapies like venetoclax. However, published *in vivo* preclinical data showed only modest survival benefit compared to saline controls (*12*), consistent with our preclinical results in Figure 6. In Phase 1 trials, [^225^Ac]-Lintuzumab was well-tolerated at doses up to 111 kBq/kg, with promising anti-leukemic activity, including significant reductions in peripheral and bone marrow blasts, and some patients achieving a morphologic leukemia-free state. However, higher doses were associated with dose-limiting toxicities such as prolonged myelosuppression, sepsis, and delayed hematopoietic recovery (*29*). While its targeted approach offers advantages over beta-emitting therapies by minimizing damage to surrounding tissues, challenges remain, including the need for better patient selection, optimizing dosing regimens, and mitigating toxicity. Another limitation of current radioligand therapies in AML is the associated damage to marrow due to higher expression of explored targets (e.g. CD33, CD123, and CLL-1) on normal myeloid cells and progenitors (*45*). As aITGB2 has undetectable expression on normal hematopoietic stem cells, targeting this integrin could avoid patient complications while still maintaining the promising anti-tumor efficacy of ^225^Ac.

We compared the therapeutic efficacy of [^225^Ac]-Macropa-PEG_4_-7065 with [^225^Ac]DOTA-antiCD33 in a pre-clinical disseminated cell line and PDX models. [^225^Ac]Macropa-PEG_4_-7065 exhibited higher therapeutic efficacy as compared to [^225^Ac]DOTA-antiCD33 with a significant improvement in survival in both studies. This improvement may be due to both the improved chelation and conjugation chemistry in [^225^Ac]Macropa-PEG_4_-7065 when compared to [^225^Ac]DOTA-antiCD33, or increased selectivity for the tumor as compared to normal tissue. PET imaging with [^89^Zr]DFO*-7065 exhibited reduced background binding to human immune cells, compared to [^89^Zr]DFO*-anti-CD33. These differences may explain the improved therapeutic efficacy of [^225^Ac]Macropa-PEG_4_-7065.

There are important limitations to this study that merit discussion, and avenues for further investigation. One potential limitation is that we used full-length IgG for evaluating therapeutic efficacy. A longer serum/biological half-life may be advantageous in the context of AML, where localization within the hematopoietic compartment is preferred, and enhanced tissue penetration is less critical. Extended exposure may also facilitate the eradication of heterogeneous leukemic cell populations. However, the risk of marrow suppression remains a concern and will require clinical evaluation (*46*). Possible avenues for future investigation of optimized therapeutic index for aITGB2-targeting RIT include comparison to antibody fragments or peptide-based conjugates to reduce systemic circulation time. Combination strategies using [^225^Ac]Macropa-PEG_4_-7065 with existing AML therapies such as hypomethylating agents (e.g., azacitidine) or BCL-2 inhibitors (e.g., venetoclax) may further enhance therapeutic efficacy. Additionally, optimization of dosing regimens, combination strategies could help maximize bone marrow targeting while minimizing off-target toxicity.

Overall, this study builds upon our prior identification of new target aITGB2 as a novel AML target through surface proteomics (*21*). These studies confirm that [^225^Ac]Macropa-PEG_4_-7065 has promising therapeutic efficacy and an improved safety profile as compared to previously reported agents, supporting its potential for clinical translation as a targeted radiopharmaceutical therapy for AML.

## Methods

### Antibody Conjugation with Chelators

The 7065 antibody (5.75 mg/mL) was prepared as previously reported (*21*). For DFO*-NCS conjugation, 173.9 μL (1 mg) of 7065 IgG in PBS was buffer-exchanged into 0.1 M Na_2_CO_3_/NaHCO_3_ (pH 9.5) using three centrifugation cycles at 10,000 rpm for 8 minutes with a YM30K MW centrifugal filter (Millipore, MA, USA), yielding a final concentration of 7.23 mg/mL. DFO*-NCS (ABX, #7272) was pre-dissolved in anhydrous DMSO (1 mg/40 μL), and 7065 was incubated with 5 equivalents (1.27 μL) at 37°C for 45 minutes. The mixture was purified using a PD10 gel column (Cytiva) pre-conditioned and eluted with 0.25 M NaOAc (pH 6). A non-targeting IgG antibody (negative control; AbCam, #AB91102) was conjugated with DFO*-NCS using the same protocol.

For conjugation with Macropa-PEG_4_-TFP, the 7065 antibody underwent buffer exchange through three cycles with 0.1 M Na_2_CO_3_/NaHCO_3_ buffer at pH 9.5 by centrifugation at 10,000 rpm for eight minutes, achieving a final concentration around 10-15 mg/mL (n=10 synthesis). The bifunctional chelator Macropa-PEG_4_-TFP was prepared as previously described (*24*). The 7065 antibody was incubated with 7.5 equivalents of Macropa-PEG_4_-TFP (pre-dissolved in DMSO; 1 mg/30 μL) at 37°C for 2 hours. Following the conjugation process, the reaction mixture was passed through a PD10 column, and elution was performed with 0.25 M NaOAc buffer. The conjugation of IgG with Macropa-PEG_4_-TFP was carried out similarly to the conjugation of the 7065 antibody. The resulting conjugate, Macropa-PEG_4_-7065 and Macropa-PEG_4_-IgG, were stored at -20°C and used without further purification for analysis and radiolabeling. Lintuzumab (anti-CD33) antibody was purchased from Creative Biolabs (catalog number TAB-756), and the conjugation of the anti-CD33 antibody with DOTA-NCS was performed according to a previously published procedure (*12*).

### Radiolabeling with ^89^Zr(C_2_O_4_)_2_, ^134^CeCl_3_ and ^225^Ac(NO_3_)_3_

For labeling with Zirconium-89, a one step process was used. ^89^Zr-oxalate was purchased from 3D Imaging (Little Rock, AR). ^89^Zr-oxalate (37 MBq, 3-5 µL) was mixed with an equal amount of 1 M Na_2_CO_3_ (3-5 µL) and 200 µL of 1M NH_4_OAc, followed by the addition of 130 µg of DFO*-7065 (7.2 mg/mL, pre-dissolved in 0.25 M NaOAc) and the mixture was incubated at 37°C for 1 hour. The mixture was subjected to instant thin-layer chromatography (iTLC) for radiochemical yield (eluted with 10 mM EDTA, pH 5.5) to confirm reaction completion, and purified by eluting through PD10 column with 0.9% saline as mobile phase. A similar procedure was followed for ^89^Zr radiolabeling of DFO*-IgG.

^134^Ce(NO_3_)_3_ in 0.1 M HCl was purchased from Department of Energy Isotope Production Program (*47*). For radiolabeling, ^134^CeCl_3_ (100 µL, 111 MBq) was mixed with 200 µL of 2M NH_4_OAc (pH 8), 200 µg of Macropa-PEG_4_-7065 (15.87 mg/mL) was added, and the reaction mixture was incubated at 37°C for 1 hour. The reaction yield was analyzed by iTLC SG (using 10 mM EDTA pH 5.5 as a mobile phase), allowing 60 minutes to pass to achieve secular equilibrium prior to plate imaging. The reaction mixture was purified by eluting through PD10 with 0.9% saline as mobile phase.

^225^Ac(NO_3_)_3_ was purchased from the Department of Energy Isotope Production Program and produced at Oak Ridge National Laboratory via the ^229^Th generator route (*48*). The Ac-225 (approximately 27.8 MBq-29.6 MBq) was dissolved in 100 µL of 0.2 M HCl. An aliquot of 4 µL (approx. 1.11 MBq of Ac-225) was added to a vial followed by 50 µL of NH_4_OAc (2M, pH 5.8) and 20 µL of L-Ascorbic acid (150 mg/mL) and 120 µg of Macropa-PEG_4_-7065 (7.6 µL). The reaction was incubated at 37°C for 30 minutes. The radiochemical yield was monitored by ITLC-SG eluted with 10 mM EDTA (pH 5.5). For purification and buffer exchange to 0.9% saline, the reaction mixture was passed through YM30K centrifugation filter with three washes and final radioimmunoconjugate purity was analyzed by ITLC-SG. For labelling of Macropa-PEG_4_-IgG, a similar procedure was followed. For radiolabeling of DOTA-anti-CD33 with ^225^Ac, an aliquot of 4 µL (approx. 1.11 MBq) of Ac-225 was added to a vial followed by 50 µL of NH_4_OAc (2M, pH 5.8) and 20 µL of L-Ascorbic acid (150 mg/mL), and then 180 µg of DOTA-anti-CD33 (19.07 µL). The reaction was incubated at 37°C for 1 hour. The radiochemical yield was determined by ITLC-SG eluted with 10 mM EDTA (pH 5.5). For purification and buffer exchange to 0.9% saline, the same process was followed as for the [^225^Ac]Macropa-PEG_4_-7065.

### Cell Binding Assay

5 million cells (Nomo-1, Nomo-1 ITGB2 KO, HL-60, THP-1, MV411) were dissolved in 100 µL of PBS (in triplicate). 1 nM (0.076 ng; 0.50 µL, 6.92 MBq) solution of [^89^Zr]DFO*-7065 (original solution concentration is 433.3 nM) was prepared in 100 µL PBS and 1% nonfat milk (20 µL, 0.1 mg/mL) was added to each vial containing cells. The cells were incubated with [^89^Zr]DFO*-7065 solution for 1 hour at 37°C. Cells were centrifuged after one hour, and the supernatant was removed. The pallet was washed with PBS twice and radioactivity bound to the cell pellet was counted using a Hidex Gamma Counter using a 480 to 558 KeV counting window, compared to standards of known radioactivity (6.92 MBq). Cell-associated activity percentage was calculated by cell pellet activity/6.92 MBq [^89^Zr]DFO*-7065 activity.

In order to test for Fc-mediated binding, a separate experiment was performed utilizing an excess of non-targeting IgG. 1 nM (0.099 ng; 0.66 µl, 6.92 MBq) solution of [^89^Zr]DFO*-7065 (original solution concentration is 333.3 nM) was prepared in 100 µl PBS and 1% nonfat milk (20 µL, 0.1 mg/mL) was added to each vial containing cells. Cells were centrifuged after one hour and the supernatant was removed. The pellet was washed with PBS twice and radioactivity bound to the cell pellet was counted using a Hidex Gamma Counter. The radioactivity associated with cell pallet and 0.5 nM (9.25 kBq solution) was counted utilizing the same method described earlier.

### Saturation Binding Assay

The k_d_ value of [^89^Zr]DFO*-7065 with aITGB2 expressing cell lines (Nomo-1, Nomo-1 ITGB2 KO, HL-60, THP-1, MV411) was determined through saturation binding assay. Aliquots of 1 million of each cell type were dissolved in 100 µL PBS, with 15 vials total for each cell line. 20 µL of 0.1% milk in PBS was added to block the non-specific binding. After 1 hour in incubation at 37°C, various concentrations of [^89^Zr]DFO*-7065 (0.001 nM-10 nM, 100 uL/vial, triplicate) in PBS was added to each vial and incubated at 37°C for 1 hour. Cells were centrifuged after one hour and supernatant was removed. The pellet was washed with 800 µL PBS twice and radioactivity bound to cell pellet was counted using a Hidex Gamma Counter. K_d_ values and B_max_ were calculated by non-linear regression, utilizing one-site specific binding using GraphPad Prism Software.

### Therapy Study in Nomo-1 disseminated model

One million Nomo-1 cells were administered by tail vein injection to NRG (NOD.Cg-*Rag1^tm1Mom^ Il2rg^tm1Wjl^*/SzJ, Stock number 007799, UCSF Mount Zion Facility) mice. Fifteen (15) days after injection of Nomo-1 cells, bioluminescence imaging was performed to monitor tumor burden. On day 16 post cell inoculation, (considered day 0 of the treatment study), the treatment study was initiated. Mice were randomized into five treatment groups (n=8) based on the average BLI signal, with a radiance range of 10^4^ to 10^6^ p/sec/cm²/sr. The treatment arms included: 1) vehicle control (100 µL saline), 2) 9.25 kBq of [^225^Ac]Macropa-PEG_4_-IgG, 3) 9.25 kBq [^225^Ac]DOTA-anti-CD33, and 4) 9.25 kBq of [^225^Ac]Macropa-PEG_4_-7065. An additional fractionated dose treatment, involving a total of three doses of 9.25 kBq of [^225^Ac]Macropa-PEG_4_-7065, was also included in the study. The fractionated doses (9.25 kBq) of [^225^Ac]Macropa-PEG_4_-7065 were administered on days 0, 28, and 56, respectively. All mice were administered 0.5 mg of cold non-targeting IgG for Fc blocking. Tumor growth in each mouse was monitored weekly using BLI. Body condition score, mobility, and body weight were monitored every other day. If body weight loss exceeded 20% or the mice exhibited deteriorating conditions, such as paralysis, hyperactivity, or head tilting, they were euthanized. After euthanasia, bone marrow and spleen were harvested and used for flow cytometry analysis of the ITGB2 population in relapsed mice after treatment. On day 150, the study was terminated.

### Statistical Significance

All the data were expressed as mean ± SD. Data was analyzed using GraphPad Prism 8 and a P value < 0.05 was considered statically significant. Two-way ANOVA was used for calculation of biodistribution values and tumor to background ratio values. The log-rank sum test was used for survival analysis.

## Data and Material Availability

The data presented in this study and in the Supplementary Information are available from the corresponding author upon request.

## Acknowledgements

R.R. Flavell and Arun P. Wiita acknowledge funding from NIH RO1 CA 297845. ^225^Ac and ^134^Ce were supplied by the U.S Department of Energy Isotope Program. NOD.Cg-Prkdc Il2rg Tg (SV40/HTLVIL3,CSF2)10-7Jic/JicTac Engrafted, CD34+ huHSCs female were provided by Taconic Biosciences. MALDI-MS data was provided by the Mass Spectrometry Facility, Department of Chemistry, University of Alberta (Edmonton, Alberta, Canada). Blood cells counts and organ functions test and histology staining procedure were performed by Comparative Pathology, UC Davis, Davis, California. The authors acknowledge the NIH grant S10 OD034286 that supported the purchase of the Mediso small animal PET/CT scanner.

## Authors’ Contributions

A. Wadhwa: Conceptualization, data curation, validation, investigation, visualization, methodology, writing-original draft, writing review and editing. H. Johnsons: Data Curation, software, visualization. K.N. Bobba: Conceptualization, validation, data curation, investigation, writing reviewing and editing. A. P. Bidkar: Conceptualization, data curation, validation, investigation, visualization, writing review and editing. E. Mayne: Conceptualization, validation, data curation, investigation, writing reviewing and editing. S. Rampersaud: Data curation, software, visualization. K. Mandal: Conceptualization, validation, data curation, investigation, writing reviewing and editing. A. S. Kang: Conceptualization, data curation, validation, investigation, visualization, writing review and editing. N. Greenland: Data curation, software, visualization and validation. R. Peter: Conceptualization, data curation, validation, investigation, visualization. A. Raveendran: Data curation, formal analysis, validation, investigation. S. Naik: Data curation, formal analysis. M. Basak: Data curation and formal analysis. A. Barpanda: Conceptualization, data curation, validation, investigation, visualization, writing review and editing. S. Prudhvi: Conceptualization, data curation, validation, investigation, visualization, writing review and editing. J. Huebner: Formal analysis, investigation, methodology. M. L. Alvarez: Conceptualization, data curation, validation, investigation, visualization, writing review and editing. S. Lee: Conceptualization, data curation, validation, investigation, visualization, writing review and editing. V. Steri: Conceptualization, data curation, validation, investigation, visualization, writing review and editing. J. J. Adams: Conceptualization, validation, data curation, investigation. S. S. Sidhu: Resources, acquisition, investigation, visualization, methodology, writing–review. D. M. Wilson: Supervision, validation, visualization, project administration, writing–review and editing. Y. Seo: Conceptualization, resources, data curation, software, supervision, validation, investigation, visualization, methodology, writing–review and editing. H. F. VanBrocklin: Data curation, software, supervision, investigation, visualization, project administration, writing–review and editing. A. C. Logan: Data curation, software, supervision, investigation, visualization, project administration, writing–review and editing. A. P. Wiita: Conceptualization, resources, data curation, software, formal analysis, supervision, funding acquisition, validation, investigation, visualization, methodology, writing–original draft, project administration, writing–review and editing. R. R. Flavell: Conceptualization, resources, data curation, software, formal analysis, supervision, funding acquisition, validation, investigation, visualization, methodology, writing–original draft, project administration, writing–review and editing.

## References

1. Key Statistics for Acute Myeloid Leukemia (AML) (available at https://www.cancer.org/cancer/types/acute-myeloid-leukemia/about/key-statistics.html).

2. C. Lai, K. Doucette, K. Norsworthy, Recent drug approvals for acute myeloid leukemia. Journal of Hematology & Oncology 12, 100 (2019).

3. G. W. Roloff, O. Odenike, A. Bajel, A. H. Wei, N. Foley, G. L. Uy, Contemporary Approach to Acute Myeloid Leukemia Therapy in 2022. Am Soc Clin Oncol Educ Book, 568–583 (2022).

4. G. Magee, B. K. Ragon, Allogeneic hematopoietic cell transplantation in acute myeloid leukemia. Best Practice & Research Clinical Haematology 36, 101466 (2023).

5. A. Ehninger, M. Kramer, C. Röllig, C. Thiede, M. Bornhäuser, M. von Bonin, M. Wermke, A. Feldmann, M. Bachmann, G. Ehninger, U. Oelschlägel, Distribution and levels of cell surface expression of CD33 and CD123 in acute myeloid leukemia. Blood Cancer J 4, e218 (2014).

6. S. Jambon, J. Sun, S. Barman, S. Muthugounder, X. R. Bito, A. Shadfar, A. E. Kovach, B. L. Wood, V. Thoppey Manoharan, A. S. Morrissy, D. Bhojwani, A. S. Wayne, M. A. Pulsipher, Y.-M. Kim, S. Asgharzadeh, C. Parekh, B. Moghimi, CD33–CD123 IF-THEN Gating Reduces Toxicity while Enhancing the Specificity and Memory Phenotype of AML-Targeting CAR-T Cells. Blood Cancer Discovery 6, 55–72 (2025).

7. A. Rondon, J. Rouanet, F. Degoul, Radioimmunotherapy in Oncology: Overview of the Last Decade Clinical Trials. Cancers (Basel) 13, 5570 (2021).

8. J. Strosberg, G. El-Haddad, E. Wolin, A. Hendifar, J. Yao, B. Chasen, E. Mittra, P. L. Kunz, M. H. Kulke, H. Jacene, D. Bushnell, T. M. O’Dorisio, R. P. Baum, H. R. Kulkarni, M. Caplin, R. Lebtahi, T. Hobday, E. Delpassand, E. V. Cutsem, A. Benson, R. Srirajaskanthan, M. Pavel, J. Mora, J. Berlin, E. Grande, N. Reed, E. Seregni, K. Öberg, M. L. Sierra, P. Santoro, T. Thevenet, J. L. Erion, P. Ruszniewski, D. Kwekkeboom, E. Krenning, Phase 3 Trial of 177Lu-Dotatate for Midgut Neuroendocrine Tumors. New England Journal of Medicine 376, 125–135 (2017).

9. O. Sartor, J. de Bono, K. N. Chi, K. Fizazi, K. Herrmann, K. Rahbar, S. T. Tagawa, L. T. Nordquist, N. Vaishampayan, G. El-Haddad, C. H. Park, T. M. Beer, A. Armour, W. J. Pérez-Contreras, M. DeSilvio, E. Kpamegan, G. Gericke, R. A. Messmann, M. J. Morris, B. J. Krause, Lutetium-177–PSMA-617 for Metastatic Castration-Resistant Prostate Cancer. New England Journal of Medicine 385, 1091–1103 (2021).

10. H. Goto, Y. Shiraishi, S. Okada, Continuing progress in radioimmunotherapy for hematologic malignancies. Blood Reviews 69, 101250 (2025).

11. T. L. Rosenblat, M. R. McDevitt, J. A. Carrasquillo, N. Pandit-Taskar, M. G. Frattini, P. G. Maslak, J. H. Park, D. Douer, D. Cicic, S. M. Larson, D. A. Scheinberg, J. G. Jurcic, Treatment of Patients with Acute Myeloid Leukemia with the Targeted Alpha-Particle Nano-Generator Actinium-225-Lintuzumab. Clin Cancer Res 28, 2030–2037 (2022).

12. R. Garg, K. J. H. Allen, W. Dawicki, E. M. Geoghegan, D. L. Ludwig, E. Dadachova, 225Ac-labeled CD33-targeting antibody reverses resistance to Bcl-2 inhibitor venetoclax in acute myeloid leukemia models. Cancer Med 10, 1128–1140 (2020).

13. S. Abedin, P. Brodin, M. Chen, M. Rotibi, U. Syed, A. Desai, E. Atallah, Safety and Dosimetric Analysis of Lintuzumab-Ac225 in Combination with Intensive CLAG-M Chemotherapy in Patients with Relapsed/Refractory AML. Journal of Nuclear Medicine 65, 241340–241340 (2024).

14. Clinical Trial: NCT03441048 - My Cancer Genome (available at https://www.mycancergenome.org/content/clinical_trials/NCT03441048/).

15. First Clinical Trial of Actimab-A Triplet Combination to Begin in AML Blood Cancers Today (available at https://www.bloodcancerstoday.com/post/first-clinical-trial-of-actimab-a-triplet-combination-to-begin-in-aml).

16. G. S. Laszlo, E. H. Estey, R. B. Walter, The past and future of CD33 as therapeutic target in acute myeloid leukemia. Blood Rev. 28, 143–153 (2014).

17. E. L. Atallah, J. J. Orozco, M. Craig, M. Y. Levy, L. E. Finn, S. S. Khan, A. E. Perl, J. H. Park, G. J. Roboz, W. Tse, K. H. Begna, R. Mawad, D. A. Rizzieri, M. S. Berger, J. G. Jurcic, A Phase 2 Study of Actinium-225 (225Ac)-Lintuzumab in Older Patients with Untreated Acute Myeloid Leukemia (AML) - Interim Analysis of 1.5 µci/Kg/Dose. Blood 132, 1457–1457 (2018).

18. R. B. Walter, Where do we stand with radioimmunotherapy for acute myeloid leukemia? Expert Opin Biol Ther 22, 555–561 (2022)

19. P. Vo, T. A. Gooley, J. G. Rajendran, D. R. Fisher, J. J. Orozco, D. J. Green, A. K. Gopal, R. Haaf, M. Nartea, R. Storb, F. R. Appelbaum, O. W. Press, J. M. Pagel, B. M. Sandmaier, Yttrium-90-labeled anti-CD45 antibody followed by a reduced-intensity hematopoietic cell transplantation for patients with relapsed/refractory leukemia or myelodysplasia. Haematologica 105, 1731–1737 (2020).

20. J. M. Pagel, F. R. Appelbaum, J. F. Eary, J. Rajendran, D. R. Fisher, T. Gooley, K. Ruffner, E. Nemecek, E. Sickle, L. Durack, J. Carreras, M. M. Horowitz, O. W. Press, A. K. Gopal, P. J. Martin, I. D. Bernstein, D. C. Matthews, 131I–anti-CD45 antibody plus busulfan and cyclophosphamide before allogeneic hematopoietic cell transplantation for treatment of acute myeloid leukemia in first remission. Blood 107, 2184–2191 (2006).

21. K. Mandal, G. Wicaksono, C. Yu, J. J. Adams, M. R. Hoopmann, W. C. Temple, A. Izgutdina, B. P. Escobar, M. Gorelik, C. H. Ihling, M. A. Nix, A. Naik, W. H. Xie, J. Hübner, L. A. Rollins, S. M. Reid, E. Ramos, C. Kasap, V. Steri, J. A. C. Serrano, F. Salangsang, P. Phojanakong, M. McMillan, V. Gavallos, A. D. Leavitt, A. C. Logan, C. M. Rooney, J. Eyquem, A. Sinz, B. J. Huang, E. Stieglitz, C. C. Smith, R. L. Moritz, S. S. Sidhu, L. Huang, A. P. Wiita, Structural surfaceomics reveals an AML-specific conformation of integrin β2 as a CAR T cellular therapy target. Nat Cancer 4, 1592–1609 (2023).

22. J. Vanhooren, R. Dobbelaere, C. Derpoorter, L. Deneweth, L. Van Camp, A. Uyttebroeck, B. De Moerloose, T. Lammens, CAR-T in the Treatment of Acute Myeloid Leukemia: Barriers and How to Overcome Them. HemaSphere 7, e937 (2023).

23. M. Ruella, J. Xu, D. M. Barrett, J. A. Fraietta, T. J. Reich, D. E. Ambrose, M. Klichinsky, O. Shestova, P. R. Patel, I. Kulikovskaya, F. Nazimuddin, V. G. Bhoj, E. J. Orlando, T. J. Fry, H. Bitter, S. L. Maude, B. L. Levine, C. L. Nobles, F. D. Bushman, R. M. Young, J. Scholler, S. I. Gill, C. H. June, S. A. Grupp, S. F. Lacey, J. J. Melenhorst, Induction of resistance to chimeric antigen receptor T cell therapy by transduction of a single leukemic B cell. Nat Med 24, 1499–1503 (2018).

24. D. J. Vugts, C. Klaver, C. Sewing, A. J. Poot, K. Adamzek, S. Huegli, C. Mari, G. W. M. Visser, I. E. Valverde, G. Gasser, T. L. Mindt, G. A. M. S. van Dongen, Comparison of the octadentate bifunctional chelator DFO*-pPhe-NCS and the clinically used hexadentate bifunctional chelator DFO-pPhe-NCS for 89Zr-immuno-PET. Eur J Nucl Med Mol Imaging 44, 286–295 (2017).

25. M. Chomet, M. Schreurs, M.J. Bolijn, M. Verlaan, W. Beaino, K. Brown, A.J. Poot, A.D. Windhorst, H. Gill, J. Marik, S. Williams, J. Cowell, G. Gasser, T. L. Mindt, G.A. M.S. Dongen, D. J. Vugts, Head-to-head comparison of DFO* and DFO chelators: selection of the best candidate for clinical 89Zr-immuno-PET. European journal of nuclear medicine and molecular imaging, 48, 694–707 (2020)

26. K. N. Bobba, A. P. Bidkar, N. Meher, C. Fong, A. Wadhwa, S. Dhrona, A. Sorlin, S. Bidlingmaier, B. Shuere, J. He, Evaluation of 134Ce/134La as a PET Imaging Theranostic Pair for 225Ac α-Radiotherapeutics. Journal of Nuclear Medicine 64, 1076–1082 (2023).

27. K. N. Bobba, A. P. Bidkar, A. Wadhwa, N. Meher, S. Drona, A. M. Sorlin, S. Bidlingmaier, L. Zhang, D. M. Wilson, E. Chan, N. Y. Greenland, R. Aggarwal, H. F. VanBrocklin, J. He, J. Chou, Y. Seo, B. Liu, R. R. Flavell, Development of CD46 targeted alpha theranostics in prostate cancer using 134Ce/225Ac-Macropa-PEG4-YS5. Theranostics 14, 1344–1360 (2024).

28. N. A. Thiele, V. Brown, J. M. Kelly, A. Amor-Coarasa, U. Jermilova, S. N. MacMillan, A. Nikolopoulou, S. Ponnala, C. F. Ramogida, A. K. H. Robertson, C. Rodríguez-Rodríguez, P. Schaffer, C. Williams, J. W. Babich, V. Radchenko, J. J. Wilson, An Eighteen-Membered Macrocyclic Ligand for Actinium-225 Targeted Alpha Therapy. Angew Chem Int Ed 56, 14712– 14717 (2017).

29. J. G. Jurcic, T. L. Rosenblat, M. R. McDevitt, N. Pandit-Taskar, J. A. Carrasquillo, S. M. Chanel, C. Ryan, M. G. Frattini, D. Cicic, S. M. Larson, D. A. Scheinberg, Phase I trial of the targeted alpha-particle nano-generator actinium-225 (225Ac-lintuzumab) (anti-CD33; HuM195) in acute myeloid leukemia (AML). JCO 29, 6516–6516 (2011).

30. E. Atallah, M. Berger, J. Jurcic, G. Roboz, W. Tse, R. Mawad, D. Rizzieri, K. Begna, J. Orozco, M. Craig, M. Y. Levy, L. Finn, K. Sharif, A. Perl, J. Park, A Phase 2 Study of Actinium-225 (225Ac)-lintuzumab in Older Patients with Untreated Acute Myeloid Leukemia (AML). Journal of Medical Imaging and Radiation Sciences 50, S109 (2019)

31. S. Y. T. Lim, F. M. Cole, G. S. Laszlo, M. C. Lunn-Halbert, J. Huo, J. Li, A. R. Kehret, R. B. Walter, Targeting the Membrane-Proximal Domain of CD33 to Maximize the Efficacy of Natural Killer Cell-Based Immunotherapies. Blood 144, 3436 (2024).

32. U. B. Hagemann, K. Wickstroem, E. Wang, A. O. Shea, K. Sponheim, J. Karlsson, R. M. Bjerke, O. B. Ryan, A. S. Cuthbertson, In Vitro and In Vivo Efficacy of a Novel CD33-Targeted Thorium-227 Conjugate for the Treatment of Acute Myeloid Leukemia. Molecular Cancer Therapeutics 15, 2422–2431 (2016).

33. J. Schwartz, J. S. Jaggi, J. A. O’Donoghue, S. Ruan, M. McDevitt, S. M. Larson, D. A. Scheinberg, J. L. Humm, Renal uptake of bismuth-213 and its contribution to kidney radiation dose following administration of actinium-225-labeled antibody. Phys Med Biol 56, 721–733 (2011).

34. F. R. Appelbaum, H. Gundacker, D. R. Head, M. L. Slovak, C. L. Willman, J. E. Godwin, J. E. Anderson, S. H. Petersdorf, Age and acute myeloid leukemia. Blood 107, 3481–3485 (2006).

35. C. Tian, Z. Chen, Immune therapy: a new therapy for acute myeloid leukemia. Blood Sci 5, 15–24 (2023).

36. L. Pérez-Amill, À. Bataller, J. Delgado, J. Esteve, M. Juan, N. Klein-González, Advancing CART therapy for acute myeloid leukemia: recent breakthroughs and strategies for future development. Front. Immunol. 14 (2023), doi:10.3389/fimmu.2023.1260470.

37. E. Atilla, K. Benabdellah, The Black Hole: CAR T Cell Therapy in AML. Cancers (Basel) 15, 2713 (2023).

38. D. Bunjes, Radioimmunotherapy of acute myeloid leukemia: a critical assessment of its prospects and limitations. Expert Rev Hematol 18, 81–89 (2025).

39. J. M. Burke, P. C. Caron, E. B. Papadopoulos, C. R. Divgi, G. Sgouros, K. S. Panageas, R. D. Finn, S. M. Larson, R. J. O’Reilly, D. A. Scheinberg, J. G. Jurcic, Cytoreduction with iodine-131-anti-CD33 antibodies before bone marrow transplantation for advanced myeloid leukemias. Bone Marrow Transplant 32, 549–556 (2003).

40. M. A. Schwartz, D. R. Lovett, A. Redner, R. D. Finn, M. C. Graham, C. R. Divgi, L. Dantis, T. S. Gee, M. Andreeff, L. J. Old, Dose-escalation trial of M195 labeled with iodine 131 for cytoreduction and marrow ablation in relapsed or refractory myeloid leukemias. J Clin Oncol 11, 294–303 (1993)

41. J. G. Jurcic, C. R. Divgi, M. R. Mcdevitt. Phase I trial of yttrium-90-HuM195 in myeloid leukemia potential for myeloablation. Cancer Biother Radiopharm. abstract 39, 15, 402–404 (2000).

42. J. G. Jurcic, Targeted Alpha-Particle Therapy for Hematologic Malignancies. J Med Imaging Radiat Sci 50, S53–S57 (2019).

43. G. Sgouros, Å. M. Ballangrud, J. G. Jurcic, M. R. McDevitt, J. L. Humm, Y. E. Erdi, B. M. Mehta, R. D. Finn, S. M. Larson, D. A. Scheinberg, Pharmacokinetics and Dosimetry of an α-Particle Emitter Labeled Antibody: 213Bi-HuM195 (Anti-CD33) in Patients with Leukemia. Journal of Nuclear Medicine 40, 1935–1946 (1999).

44. J. G. Jurcic, T. L. Rosenblat, Targeted Alpha-Particle Immunotherapy for Acute Myeloid Leukemia. Am Soc Clin Oncol Educ Book, e126–e131 (2014).

45. A. Ehninger, M. Kramer, C. Röllig, C. Thiede, M. Bornhäuser, M. von Bonin, M. Wermke, A. Feldmann, M. Bachmann, G. Ehninger, U. Oelschlägel, Distribution and levels of cell surface expression of CD33 and CD123 in acute myeloid leukemia. Blood Cancer J 4, e218 (2014).

46. C. D. DiNardo, A. H. Wei, How I treat acute myeloid leukemia in the era of new drugs. Blood 135, 85–96 (2020).

47. T. A. Bailey, V. Mocko, K. M. Shield, D. D. An, A. C. Akin, E. R. Birnbaum, M. Brugh, J. C. Cooley, J. W. Engle, J. W. Fassbender, S. S. Gauny, A. L. Lakes, F. M. Nortier, E. M. O’Brien, F. D. White, C. Vermeulen, S. A. Korimor, R. J. Abergel, Developing the 134Ce and 134La pair as companion positron emission tomography diagnostic isotopes for 225Ac and 227Th radiotherapeutics. Nature chemistry 13, 284–289 (2021)

48. S. van Cleve, R. Boll, P. Benny, T. Dyke, J. Kehn, K. Phillips, Thorium-229 Generator Production of Actinium-225 at Oak Ridge National Laboratory. Journal of Medical Imaging and Radiation Sciences 50, S11–S12 (2019).

