## Supporting information for "Effective imaging and treatment of Acute Myeloid Leukemia with radiotheranostics targeting the activated conformation of integrin-βeta2"

.

*** Corresponding authors:**

*Arun P. Wiita,;

1450 Third Street, HD265, Helen Diller Cancer Research Building,

*Robert R. Flavell,;

185 Berry St., Lobby 6, Suite 350

**Methods**

**General**

The 7065-antibody was generated and purified as described previously *(1)*. DFO*-NCS was purchased from ABX (Catalog No. 7272) and p-isothiocyanate-benzyl-DOTA (Catalog No. B-205) were purchased from Macrocyclics. Native human IgG (Catalog No. ab91102) was purchased from Abcam. iTLC-SG (Catalog No. SG10001) was purchased from Agilent Technologies.

**AML Primary Sample Analysis**

Primary samples were sourced from the Hematologic Malignancies Tissue Bank at UCSF, collected under Institutional Review Board-approved protocols. Freshly acquired peripheral blood or bone marrow samples underwent red blood cell lysis using 1X ammonium chloride lysis buffer, filtration through a 70 µm filter, and a wash with flow cytometry buffer. Cryopreserved samples were rapidly thawed at 37ºC with added 10 ng/mL DNase I, followed by 70 µm filtration and flow cytometry buffer wash. Samples viability and cell counts were then taken before staining for flow cytometry analysis

**Stability Studies**

For stability studies, [^89^Zr]DFO*-7065 was purified using a PD10 pre-treated with 0.1 % ascorbic acid in saline to prevent radiolysis and reaction mixture was passed through this pre-treated PD10 column using the 0.1 % ascorbic acid as an eluent. [^89^Zr]DFO*-7065 (37 MBq radiochemical purity > 98%, 90-100 µL) was mixed with 500 µL of either saline or human serum at 37°C. At the indicated time points, 10 µL aliquots were analyzed by ITLC-SG using 10 mM EDTA (pH=5.5) as a mobile phase.

[^225^Ac]Macropa-PEG_4_-7065 (0.37 MBq-0.55 MBq, Radiochemical purity > 95 % after purification through centrifugal filtration) was diluted with 500 µL of either saline or human serum at 37°C. At the indicated time points, 5-10 µL aliquots were analyzed by ITLC-SG using 10 mM EDTA (pH 5.5) as an eluent. All radio-ITLC’s were scanned after 24 h, to achieve secular equilibrium.

**Flow Cytometry**

Flow Cytometry was performed to analyze the aITGB2+ in AML cells. Immunostaining of cells was performed as per the instructions from the antibody vendor unless stated otherwise. 1 × 10^6^ cells were resuspended in 100 µL of flow cytometry buffer (PBS + 2% FBS + 2mM EDTA) with 5µg of human Fc Block (Biolegend, 422302) added. Cells were incubated at 22 °C for 10–15 min, then 3µg of antibody was added. Cells were incubated at 4°C for 30-45 minutes protected from light then washed three times with flow cytometry buffer. For staining the active form of integrin β2, an additional antibody incubation step was performed at 37 °C for 1 hour, protected from light, before proceeding with the regular antibody staining. For staining for activated integrin β2, the flow cytometry buffer was RPMI-1640 + 5% FBS + 2% bovine serum albumin (BSA) + 50 µg mL–1 DNase I (Gold Biotechnology, D-301-500). Compensation used UltraComp eBeads Compensation Beads (Invitrogen, 01-2222-42). All samples were gated on FSC-A/SSC-A for lymphocyte population, then single cells gated in SSC-A/SSC-H, and tumor cells were gated on CD45+CD33+. Gates were determined via unstained and FMO controls. Flow cytometry was performed on the CytoFLEX platform (Beckman Coulter), and data was analyzed using FlowJo_v10.10.0.

**Cell Culture**

Nomo-1, HL-60, THP-1, MV411 cell lines were purchased from ATCC. All cells were maintained in RPMI1640 medium supplemented with 10% FBS, 100 U penicillin, and 100 µg/mL streptomycin in a humidified incubator at 37°C and 5% CO_2_. Cell lines used in our studies were negative from mycoplasma contamination when routinely tested with the bioluminescence based MycoAlert Mycoplasma Detection Kit.

**Cas9 knockouts of Nomo-1 cells**

Knockout cell lines were generated using in vitro nucleofection of Cas9 ribonuclease protein complex. Unless otherwise specified, 2 μL of each sgRNA (100 µM; Synthego Corporation) and recombinant Cas9 protein (40 µM; QB3 MacroLab, University of California, Berkeley) was incubated at 37 °C for 15 min to generate ribonuclease complex, which was then nucleofected using a 4D-Nucleofector (Lonza) with the built-in program DS-137 for cell lines (using Lonza V4XC-2032) unless otherwise specified. Knockout cell lines were allowed to recover before being purified through Fluorescent Activated Cell Sorting (Flow Cytometry). sgRNA sequences were obtained from Brunello DNA Library.

| sgRNA Target (Brunello Library) | Strand | DNA Sequence |
| --- | --- | --- |
| ItgB2 (g1) | antisense | TCAGATAGTACAGGTCGATG |
| ItgB2 (g2) | sense | CTCCAACCAGTTTCAGACCG |
| ItgB2 (g3) | antisense | TCAGGGTGCGTGTTCACGAA |
| ItgB2 (g4) | sense | TCATCCCCAAGTCAGCCGTG |

**Generation of Luc+ cells**

All cell lines Nomo-1, HL-60, THP-1, MV411, Nomo-1 ITGB2 KO cells were all modified to express mCherry and luciferase according to a previous established procedure *(2)* and transduced with a lentiviral expressing mCherry fluorescent and luciferase tag.

**Membrane Bound and Internalized assay**

3 million Nomo-1 or HL-60 cells were dissolved in 100 µL of PBS (in triplicate). 1 nM (0.099 ng; 0.66 µL) solution of [^89^Zr]DFO*-7065 (original solution concentration is 433.3 nM) was prepared in 100 µL PBS and 1% nonfat milk (20 µl, 0.1 mg/mL) and added to each vial containing cells. The total volume of the solution was 220 µL. The cells were incubated with [^89^Zr]DFO*-7065 solution for 1 h, 4 h or 24 h at 37°C. After incubation, the cells were centrifuged and washed with PBS twice. Cells were treated with a solution of 50 mM glycine/100 mM NaCl for 5 minutes at 4°C and supernatant corresponding to the membrane bound fraction was removed and separated from the pellet which contained the internalized fraction. The membrane bound and internalized fractions were calculated by activity associated with supernatant/6.92 MBq solution of 1 nM [^89^Zr]DFO*-7065.

**Immunoreactivity (Lindmo) Assay**

In centrifuge tubes, six aliquots of 5 × 10^6^ Nomo-1 cells in 500 µL PBS were prepared (n =3 for both experiment and blocking group). Additional similar aliquots were prepared containing 2.5 × 10^6^ cells, 1.25 × 10^6^ cells, 0.0625 × 10^6^ cells, 0.3125 × 10^6^ and 0.1525 × 10^6^ cells. 50 µg of cold 7065 IgG was added to the aliquots in the blocking group to saturate the antigen on the cells. All the samples were incubated on the ice for 30 min and manually tubes were agitated after every 10 mins to prevent the formation of cell pallet.

In each tube 11.9 µL of a solution containing 40 ng/mL (266.6 pM) of [^89^Zr]DFO*-7065 in PBS was added and incubated on ice 1 h. After 1 hour, all the treated and blocked samples were centrifuged at 650 rpm for 3 min and supernatant was collected in different tubes. Cells were washed twice with PBS and cell pellet and supernatant activity was counted using Hidex gamma counter. The ratio of cell associated activity to the total radioactivity of the each sample was plotted as a function of cells increasing concentration and inverse of Y-slope (1/B_max_) of the resulting curve denotes the immunoreactive fraction of the [^89^Zr]DFO*-7065.

**In vivo studies**

**General mouse methods**

All animal studies were performed according to Institutional Animal Care and Use Committed-approved protocols at the University of California (AN194778). For imaging purpose, a mixture of male and female NOD SCID gamma (commonly named as NSG, stock number 005557) aged 7 to 9 weeks old were purchased from Jackson laboratory**.** For therapy in AML disseminated model, a mixture of male and female NOD Rag gamma aged 7 to 9 weeks (commonly known as NRG, stock number from Jackson Lab 007799) were obtained from a breeding colony maintained at UCSF. NOD.Cg-Prkdc Il2rg Tg (SV40/HTLV-IL3.CSF2)10-7lic/licTac Engrafted, CD34+ huHSCs female humanized mice (HSCCB-13395-F) were purchase from Taconic Biosciences.

**In vivo disseminated model formation**

To generate the disseminated AML models, 4 million Nomo-1, Nomo-1 ITGB2 KO, HL-60, MV411, THP-1, or HL-60 cells in PBS were injected by tail vein into a cohort of mix of male and female NSG mice aged between 7 to 9 weeks. Bioluminescence Imaging was used to monitor the tumor engraftment. Tumor growth was heterogeneous and differed between cell lines, but mostly accumulated in the bone marrow and liver. When mice showed bioluminescence signal within a reference range of 10^5^ to 10^7^ photons/sec/cm^2^/steradian, mice were injected with [^89^Zr]DFO*-7065 followed by PET imaging and biodistribution study.

**Bioluminescence Imaging**

In vivo BLI was performed to monitor the tumor burden/lesions in mice inoculated with luciferase-tagged AML cells. D-Luciferin was purchased from LUCK-1G-Gold Biotechnology. One gram of D-Luciferin was dissolved in 33 mL of PBS. 100 µL solution (approx. 150 mg/kg) of Luciferin in PBS was injected intraperitoneally into mice, and mice were allowed to move freely for 8-10 minutes. After 8 minutes, mice were anesthetized and imaged using IVIS 50 (PerkinElmer). Images were acquired after 60 seconds of exposure time and the intensity of Luciferase-tagged AML cells was quantified as radiance which has a unit photons/sec/cm^2^/steradian using spherical region of interest using Living Image 4.0 software.

For ex vivo BLI, mice were injected 100 µL of D-Luciferin. After 8 minutes mice were sacrificed, organs were dissected and placed in a petri dish. All the organs are imaged with 60 seconds of exposure time and intensity of signal has been recorded in photons/sec/cm^2^/steradian.

**In vivo PET Imaging**

**General PET Imaging method**

Approximately 5 to 6 weeks after tumor implantation, when mice showed the BLI signal within a reference range 10^5^ to 10^7^ photons/sec/cm^2^/steradian, each mouse was injected with 3.7-5.55 MBq, 10 µg (n=4) of the indicated radiopharmaceutical ([^89^Zr]DFO*-7065, [^89^Zr]DFO*-IgG, [^89^Zr]DFO*-7065 + 25 fold excess of 7065, or [^134^Ce]Macropa-PEG_4_-7065). In all cases, 0.5 mg non-specific binding cold IgG was co-administered to block the Fc receptor in NSG mice (3). The animals were imaged at day 1, day 2, and day 4 post injection of all the radiotracer. µPET/CT imaging data were acquired using a small animal PET/CT scanner (nanoScan PET123S/CT1512, Mediso Medical Imaging Solutions) using a multi-animal bed that enables scanning four mice simultaneously. The mice were anesthetized with ~2% isoflurane. A custom-made intravenous catheter was used for tail vein administration of radiotracer. PET data with 15 cm axial field of view (FOV) were acquired for 20 minutes in list mode, followed by helical CT for anatomical localization and correction for attenuation and scatter. The helical pitch for CT was 1.0 for 2.57 rotations to cover the matched axial FOV of PET. X-ray tube settings were 50 kVp and 0.98 mA with exposure time of 170 ms at each angle for 360 projections per rotation. CT data were reconstructed using the vendor-provided cone-beam filtered back projection (FBP) with a cosine filter. The reconstructed CT volume was in the matrix of 486´486´603 with the isotropic voxel size of 0.25 mm. PET data were reconstructed using the vendor-provided 3D iterative algorithm with 4 iterations and 6 subsets. All corrections (randoms, attenuation, and scatter) were applied for PET reconstruction, and the vendor-developed body-air segmentation method using coregistered CT was used for attenuation correction. The reconstructed PET volume was in the matrix of 225´225´366 with the isotropic voxel size of 0.4 mm.

For [^18^F]-FDG imaging in AML models Nomo-1 and THP-1 when BLI signal reached in the reference range, mice were injected with 7.4 MBq- 8.14 MBq of [^18^F]-FDG (n=4)/group and similar imaging procedure has been followed as previously described followed by a biodistribution study *(4)*.

**Ex vivo Biodistribution and Ex vivo PET/CT imaging**

Following PET/CT imaging of mice injected with [^89^Zr]DFO*-7065 in Nomo-1, MV411, THP-1, HL-60 model and Nomo-1 ITGB2 knockout model, mice were injected with 100 µL of D-Luciferin and BLI images were acquired with 60 seconds of exposure time. Mice were sacrificed after BLI and blood was collected by cardiac puncture. Major organs (femur, liver, heart, kidney, small intestine, large intestine, spleen, pancreas, lungs, stomach and brain) were harvested and collected in a Petri dish. Organs were imaged with BLI followed by PET/CT imaging using the imaging acquisition parameters after above. After PET/CT imaging, bone marrow was extracted using a previously published procedure *(4)*. All the organs were collected in tubes and counted in Hidex Gamma Counter and percentage injected activity per gram of the tissue (%IA/g) was calculated with standards of known radioactivity.

**Size Exclusion Chromatography and optimization of Size exclusion chromatography (SEC) for [^225^Ac]Macropa-PEG_4_-7065**

The radiopharmaceuticals [^89^Zr]DFO*-7065 and [^225^Ac]Macropa-PEG_4_-7065 were analyzed using size exclusion chromatography. The LabLogic Logi-CHROM HPLC instrument was used, and the column used was a BioSep™ SEC-s3000 290 Å with dimensions of 300 × 7.8 mm.

For size exclusion chromatography analysis of [^225^Ac]Macropa-PEG_4_-7065**,** 10 µL of [^225^Ac]Macropa-PEG_4_-7065 was mixed with 40 µL of a saline and injected in Lablogic HPLC instrument and passed through 3000 SEC column. 0.1% IX PBS was used as the eluent (mobile phase) and 20 minutes of chromatogram was recorded with a flow rate of 1mL/minute. 1 mL fractions were collected tube, and the first 20 fractions were counted on Hidex after secular equilibrium and the graph between time and Count per minute was plotted.

**Cell killing Assay**

For each indicated cell line, 2000 cells were plated in a 96 well black opaque plate. Various concentration of [^225^Ac]Macropa-PEG_4_-7065, [^225^Ac]DOTA-anti-CD33 (ranging from 0.001 pCi/mL to 10 nCi/mL) in 10% RPMI media were incubated with the cells for 96 hours. After treatment, bioluminescence signals of the cells were recorded with multiplate reader (TECAN software) after 10 minutes incubation by Cell Titre Glo® according to the manufacturer’s instructions. The % cell viability was fitted into a sigmoidal dose-response curve to determine IC_50_ value using GraphPad Prism.

**Colony formation assay**

200 Nomo-1, Nomo-1 ITGB2 KO, HL-60, or MV411 cells were seeded in 1 mL of media per well in a 6-well plate. The cells were treated with varying concentrations ranging from 0.1 nCi/mL to 50 nCi/mL in 10% RPMI media of [^225^Ac]Macropa-PEG_4_-7065 for 96 hours. After treatment, the contents of each well were transferred to 1.5 mL centrifuge tubes, centrifuged, and the supernatant was discarded. The cell pellets were washed twice with PBS and subsequently resuspended in 1.5 mL of MethoCult media (H4230 without cytokines) in 6-well plates. MethoCult (#04230) was purchased from Stem Cell Technologies and was thawed overnight at 4°C or for 2–3 hours at room temperature. AML cells, being non-adherent, do not attach to surfaces; MethoCult provides a supportive medium for colony formation. The cells were incubated for 14 days at 37°C to allow colonies to form. On day 14, the colonies were counted using a microscope.

**DNA damage analysis after treatment of [^225^Ac]Macropa-PEG_4_-7065**

Nomo-1 cells were seeded in multi chamber slides. The treatment of [^225^Ac]Macropa-PEG_4_-7065 was given with a radioactivity dose of 0.0037 to 3.7 kBq/mL. After 96 hours of treatment, cells were washed with PBS and fixed with 4% formaldehyde for 10 minutes. Fixed cells were permeabilized with 1% TBST, followed by blocking with 2% nonfat milk in PBS. The cells were subsequently stained with primary antibody for phospho-γ-H2AX (1:300 dilution, anti-γ H2AX, phospho S139 antibody, Abcam #ab81299, RRID: AB_1640564) and the secondary antibody [1:1,000 dilution, anti-rabbit IgG, F(ab0)2 Fragment, Alexa Fluor 647 conjugate, #4414, RRID: AB_10693544]. The nucleus was stained with 4’,6-diamidino-2-phenylindole (DAPI; 1:1,000 dilution from 1 mg/mL stock). The images were captured with a ZEISS LSM 780 confocal microscope. The γ-H2AX foci were quantified using the Cell Profiler program, in which nuclei were first identified using DAPI fluorescence, and the H2AX foci within the nuclear regions were counted. Three images from each treatment, each containing at least six cell nuclei, were analyzed.

**Dosimetry calculation**

Dosimetry calculations were performed by determining time-integrated activity coefficients and applying curve-fitting techniques within the EXM module of OLINDA/EXM Version 1.1 *(5)*. For this analysis, the digital mouse phantom provided in OLINDA Version 2.0 was used. Biodistribution studies were performed at day 1, day 2, day 4 and day 7 using the same procedure as above to calculate % IA/gram of [^225^Ac]Macropa-PEG_4_-7065 in NRG mice bearing Nomo-1 disseminated xenograft (n=4 per group). Data from biodistribution study were systematically organized to calculate the time and percentage of injected activity for each organ and tumor. These values served as inputs to obtain time-integrated activity coefficients. Finally, the equivalent dose (Sv) was calculated by multiplying the absorbed dose (Gy) with radiation weighting factors.

Flow cytometry **study to analyze the aITGB2+ tumor population in relapsed mice**

Spleens and bone marrows were harvested from mice at humane endpoints. Samples were spun down at 400 g for 5 minutes and resuspended in 1X Ammonium Chloride (ACK) Lysis, then gently mixed on a rocker at room temperature for 10 minutes. Samples were then spun at 500 g for 5 minutes and ACK lysis was decanted into ethanol. ACK Lysis steps were repeated until minimal RBC was visible in the cell pellet. Cell pellets were then resuspended in 5mL of Bulk Lyse Wash Solution (PBS + 0.1% Sodium Azide + 0.5% Bovine Serum Album (BSA)) and run through a 70 µm filter. Cells were resuspended in the flow cytometry Buffer (PBS + 2% FBS + 2mM EDTA) and proceeded to flow cytometry staining protocol. Immunostaining of cells was performed as per the instructions from the antibody vendor unless stated otherwise. Cells were resuspended at 1 × 10^6^ cells/100 µL of flow cytometry buffer with 5 µg of human Fc Block (Biolegend, 422302) added. Cells were incubated at 22 °C for 10–15 min, then 3 µg of CD45-APC (Biolegend 368512) antibody was added. Cells were incubated at 4°C for 30-45 minutes protected from light then washed three times with flow cytometry buffer and filtered through a 70 µm filter. All samples were gated on FSC-A/SSC-A for lymphocyte population, then single cells gated in SSC-A/SSC-H, and tumor cells were gated on CD45+. Cells were sorted on a BD Biosciences flow cytometry Aria Fusion 2 Cell Sorter using FMO and unstained controls to establish gates.

**Toxicity study in healthy Black C57BL/6 and NRG mice**

**Acute toxicity in black mice**

The toxicity of [^225^Ac]Macropa-PEG_4_-7065 was evaluated in healthy Black C57BL/6 mice aged 5-6 weeks (Jackson Laboratory). For acute toxicity, the mice were divided into four groups (n=5 mice per group). Treatment groups included saline control, 9.25 kBq dose of [^225^Ac]DOTA-anti-CD33, 9.25 kBq of [^225^Ac]Macropa-PEG_4_-7065, and 9.25 kBq of [^225^Ac]Macropa-PEG_4_-IgG. The mice were monitored, and their body condition score and body weights are recorded at every alternate day. At day 7, peripheral blood was withdrawn by submandibular vein and blood parameters was analyzed by Hemavet 950/950FS Multi-Species Analyzer. At day 28, mice were sacrificed, blood was collected through cardiac puncture and stored in EDTA coated tubes to prevent coagulation and blood parameters were analyzed. Serum samples were obtained by allowed the blood containing vials to sit at 4°C for 30 min to separate the serum from the clotted blood. The serum samples were separated from the clots by centrifugation at 10,000 rpm for 10 minutes at 4°C. The blood and serum samples were sent to the pathology laboratory at Comparative Pathology Laboratory, School of Veterinary Medicine, University of California Davis for analysis where blood cell counts, and organ function testing was performed.

**Chronic Toxicity in healthy NRG mice**

The toxicity of [^225^Ac]Macropa-PEG_4_-7065 was evaluated in NRG mice aged 5-6 weeks. For the chronic toxicity, the mice were divided into five groups (n=5 mice per group). Cohorts included saline, 4.62 kBq single dose, 4.62 kBq µCi fractionated dose, 9.25 kBq single dose and 9.25 kBq fractionated dose. Fractionated doses were injected every four weeks (day 0, day 28 and day 56). Body condition score and body weights are recorded at every alternate day till day 100. At day 100 mice were sacrificed, and blood parameters and organ function test were performed as the procedure described previously.

**Histopathology and Necropsy Analysis**

After euthanasia for toxicity evaluation for both acute as well as chronic toxicity, dose limiting organs including kidney, liver, lungs, spleen, heart and bone were extracted and fixed in 10% formalin (for both Acute and Chronic toxicity). Histologic analysis was carried out to examine the microscopic features of the tissues. For Hematoxylin and Eosin (H&E) staining of dose-limiting organs, the tissues were fixed in formalin, processed through EtOH gradient (30% to 70%), and embedded in paraffin. Tissue sections with a thickness of 4 µm were prepared for H & E staining at University of California Davis.

**Therapy study in disseminated Patient Derived Xenograft model**

NSG-SGM3 humanized mice (stock number 703321) were purchased from Jackson Laboratory. Mice were pre-conditioned with 6.25 mg/kg dose of busulfan (intravenously) for two days. One day after pre-conditioning NSG-SGM3 mice were injected intravenously with 1 million PDX B cells *(1)*. Following 10 days after inoculation with PDX cells, peripheral blood was withdrawn, and flow analysis was performed to monitor the PDX engraftment in the animal’s blood. On day 11 after inoculation, mice were randomized in three groups (n=5 per group). Treatment arms included 4.62 kBq dose of [^225^Ac]Macropa-PEG_4_-7065, 4.62 kBq dose of [^225^Ac]DOTA-anti-CD33, and saline vehicle control. The body condition score, mobility and body weight were recorded at every alternate day. Tumor burden was monitored by collecting peripheral blood via the submandibular vein, followed by flow cytometric analysis to quantify human CD45⁺ cells. Mice were euthanized when the hCD45⁺ cell population reached approximately 50–90% and when clinical symptoms such as reduced mobility or hunching were observed.

**Proteomic sample preparation and LC-MS/MS analysis**

The cells were washed twice with 1X PBS, followed by the addition of 500 µL of 8 M urea buffer supplemented with 1X protease inhibitor cocktail (Thermo Fisher Scientific, 1861280). Cells were then lysed via sonication, followed by a high-speed spin at 17,000 × g for 10 minutes at 4 °C to clear debris and isolate crude proteomic contents. The samples were quantified using the BCA assay, and 100 µg of protein were resuspended in Lyse (PreOmics, P.O.00027), followed by the addition of trypsin/lys-C mix for digestion as per the manufacturer’s protocol. The resulting peptide mixture was then desalted, eluted, and dried completely using a vacuum concentrator (Labconco, 7810010). Dried peptides were reconstituted in LC-load buffer, quantified, and 500 ng of peptides were loaded onto the mass spectrometer. Samples were analyzed using a Thermo Scientific Vanquish Neo liquid chromatography system coupled to an Orbitrap Eclipse mass spectrometer. Chromatographic separation was performed over a 60-minute linear gradient at a flow rate of 300 nL/min on a 60 cm column with a 75 µm inner diameter. The mass spectrometer was operated in data-independent acquisition (DIA) mode. Full MS scans were acquired at a resolution of 60,000 with a scan range of 350 to 900 m/z, and an automatic gain control (AGC) target of 300% with automatic maximum injection time. For the DIA master scan, HCD fragmentation was employed with a normalized collision energy (NCE) of 27%. DIA isolation windows of 12 m/z were used, covering a scan range of 200–1500 m/z, the same as the full MS scan. MS/MS spectra were acquired at a resolution of 30,000 *(6)*. The raw LFQ spectral data were analyzed using DIA-NN, a neural network-based tool *(7)*. The data were searched against the human-reviewed UniProt database. Trypsin was specified as the protease, allowing for up to one missed cleavage. Methionine oxidation and N-terminal acetylation were set as variable modifications, while cysteine carbamidomethylation was fixed. The analysis was conducted in library-free mode, with DIA-NN generating in silico spectral libraries directly from the protein sequence database. Default settings for precursor charge states and ion types were used for protein quantification. The LFQ intensities were median normalized and subsequently log-transformed. Differential expression of proteins was assessed using Welch’s t-test, with significance defined by a p-value < 0.05 and a log2 fold change (Log2FC) threshold of greater than 2 or less than -2 to identify significantly upregulated or downregulated proteins *(8)*.

**SUPPLEMENTARY FIGURES**

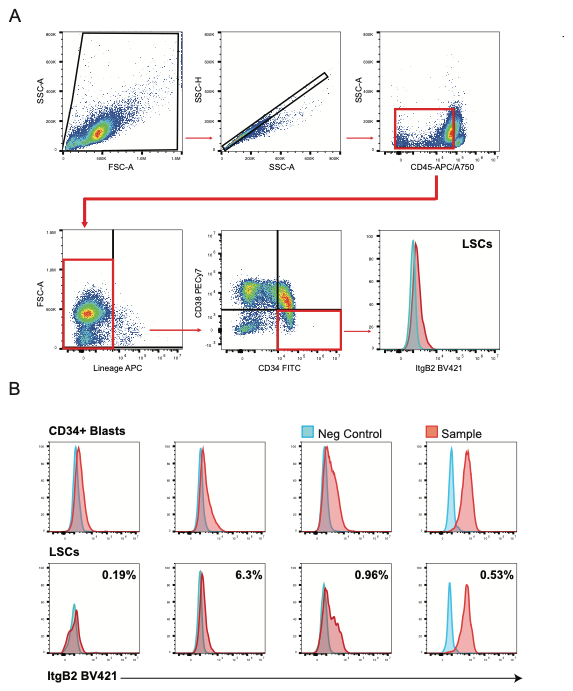
**Figure S1** Flow cytometric staining of ItgB2 on 4 thawed AML patient tumor samples selected as CD45 dim/CD34+ blasts (top row) or CD45 dim/CD34+/CD38-/Lin- (bottom row). Fluorescence minus one (FMO) used as negative gate. Each sample was measured once due to sample input limitation

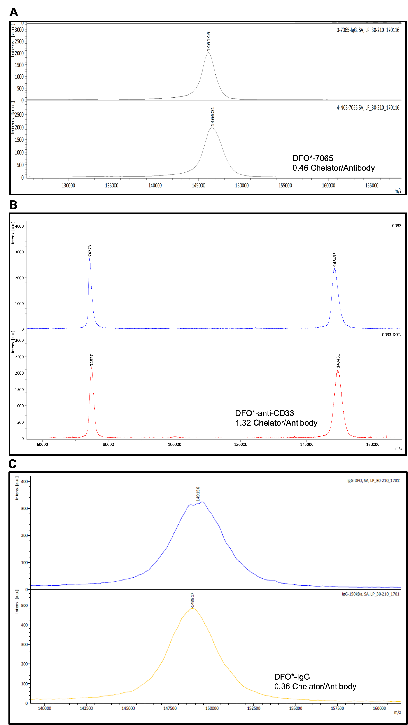

**Figure S2 MALDI-TOF mass analysis for the immunoconjugates**-Number of chelators attached per antibody was calculated by mass difference between antibody and chelator-antibody conjugate divided by the mass of the chelator A) 7065 and DFO*-7065 chromatograms. B) anti-CD33 and DFO*-anti-CD33 chromatograms. C) IgG and DFO*-IgG chromatograms.

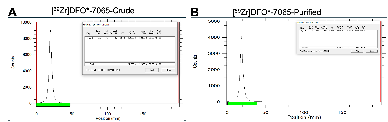

**Figure S3 ITLC of radio immunoconjugate before (Crude mixture) and after purification** Radio ITLC-SG for [^89^Zr]DFO*-7065 A) before and B) after purification.

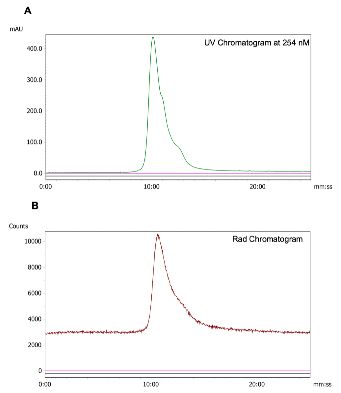

**Figure S4 HPLC chromatogram for [^89^Zr]DFO*-7065 after purification through PD10** A) UV chromatogram for [^89^Zr]DFO*-7065 at 254 nM wavelength B) corresponding Rad chromatogram for [^89^Zr]DFO*-7065

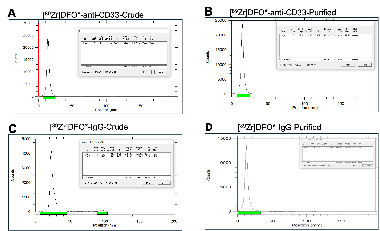

**Figure S5 ITLC of radio immunoconjugates before (Crude mixture) and after purification** Radio ITLC-SG for [^89^Zr]DFO*-anti-CD33 A) before and B) after purification. ITLC for [^89^Zr]DFO*-IgG C) before and D) after purification.

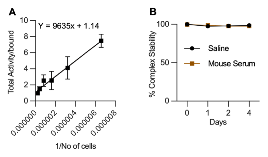

**Figure S6 LINDMO assay and stability assessment of [^89^Zr]DFO*-7065 in saline and mouse serum** A) [^89^Zr]DFO*-7065 demonstrated an immunoreactive fraction of 87.8 ± 0.65% with Nomo-1 cells expressing aITGB2 B) [^89^Zr]DFO*-7065 showed over 99% of the radioimmunoconjugate stability in both saline and human serum for seven days at 37°C.

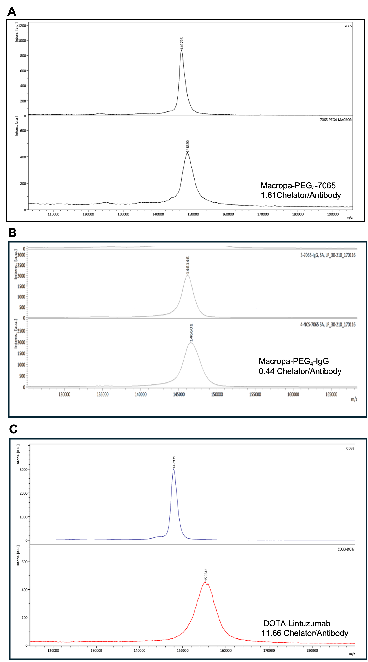

**Figure S7 MALDI-TOF mass analysis for the immunoconjugates**-Number of chelators attached per antibody was calculated by mass difference between antibody and chelator-antibody conjugate divided by the mass of the chelator A) 7065 and Macropa-PEG_4_-7065 chromatograms. B) IgG and Macropa-PEG_4_-IgG chromatograms. C) anti-CD33 and DOTA-anti-CD33 chromatograms.

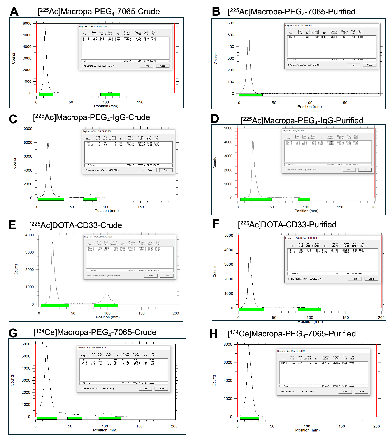

**Figure S8 ITLC of radio immunoconjugates before (Crude mixture) and after purification** Radio ITLC-SG for [^225^Ac]Macropa-PEG_4_-7065 A) before and B) after purification. Radio ITLC-SG for [^225^Ac]Macropa-PEG_4_-IgG C) before and D) after purification. ITLC for [^225^Ac]DOTA-anti-CD33 E) before and F) after purification. Radio ITLC-SG for [^134^Ce]Macropa-PEG_4_-7065 G) before and H) after purification.

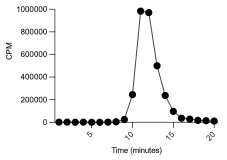

**Figure S9 HPLC rad chromatogram for [^225^Ac]Macropa-PEG_4_-7065.** Size exclusion chromatography was performed following injection of [^225^Ac]Macropa-PEG_4_-7065. One mL fraction of [^225^Ac]Macropa-PEG_4_-7065 was collected per tube (20 fractions total), and were counted on Hidex after secular equilibrium.

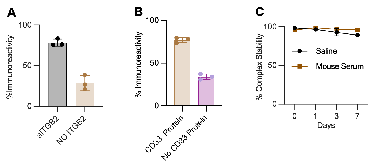

**Figure S10 [^225^Ac]Macropa-PEG_4_-7065 demonstrates high specificity towards aITGB2** A) Magnetic bead assay with after incubation with [^225^Ac]Macropa-PEG_4_-7065 with recombinant ITGB2 protein showed immunoreactive fraction of 78.64 ± 4.28 %. which is reduced to 28.28 ± 8.94 % in absence of recombinant ITGB2 protein. B) Magnetic bead assay with [^225^Ac]DOTA-anti-CD33 with recombinant CD33 protein showed immunoreactive fraction of 77.04 ± 3.15%. which is reduced to 34.07 ± 3.19 % in absence of recombinant CD33 protein. C) [^225^Ac]Macropa-PEG_4_-7065 showed over 99% of the radioimmunoconjugate stability in both saline and human serum for seven days at 37°C.

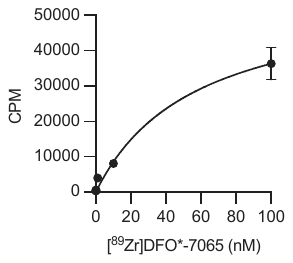

K_d_ =56.46 ± 1.82 nM

**Figure S11** K_d_ measurement of [^89^Zr]DFO*-7065 on MV411 cell line determined by a saturation binding assay (K_d_ = 56.46 ± 1.82 nM)

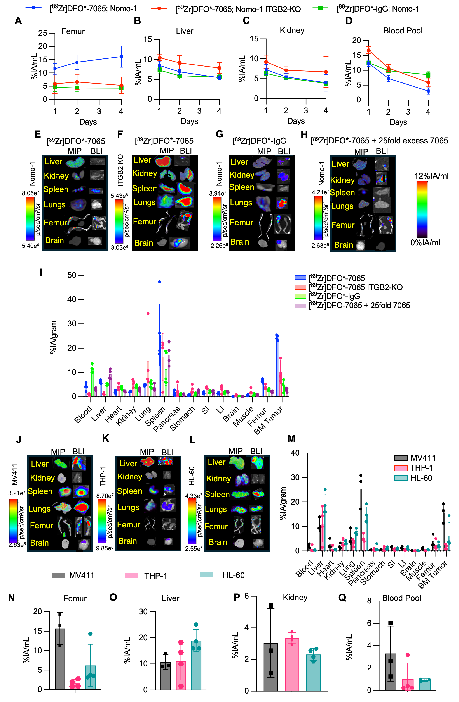

**Figure S12 [^89^Zr]DFO*-7065 PET can detect AML disseminated lesions.** Region of Interest (ROI) graph indicating % ID/gram accumulation of [^89^Zr]DFO*-7065 in Nomo-1 Luc model (blue color), [^89^Zr]DFO*-7065 in Nomo-1 ITGB2 KO (red color) and [^89^Zr]DFO*-IgG in Nomo-1 Luc model (green color) at 1day, 2days, 4days post injection in A) Femur B) Liver C) Kidney D) Cardiac blood Pool. Ex vivo PET/CT and BLI images of E) [^89^Zr]DFO*-7065 in Nomo-1-Luc model F) [^89^Zr]DFO*-7065 in Nomo-1 ITGB2 KO model G) [^89^Zr]DFO*-IgG in Nomo-1 model H) [^89^Zr]DFO*-7065 + 25 fold excess of 7065 in Nomo-1 Luc model I) Ex vivo biodistribution of [^89^Zr]DFO*-7065 in Nomo-1 Luc model, [^89^Zr]DFO*-7065 in Nomo-1 ITGB2 KO model, [^89^Zr]DFO*-IgG in Nomo-1 Luc model, [^89^Zr]DFO*-7065 + 25 fold excess of 7065 in NSG mouse at 4 days post injection. Ex vivo PET/CT and BLI images of [^89^Zr]DFO*-7065 in J) MV411-Luc model K) THP-1-Luc model L) HL-60-Luc model M) Ex vivo biodistribution indicating %IA/gram accumulation of [^89^Zr]DFO*-7065 in different organs in MV411-Luc, HL-60-Luc, THP-1-Luc model at 4 days post injection. Region of Interest (ROI) graph indicating % IA/gram accumulation of [^89^Zr]DFO*-7065 in MV411-Luc (black color), HL-60-Luc (pink color), THP-1 (green color) respectively at 4 days post injection in N) Femur O) Liver P) Kidney and Q) Blood pool.

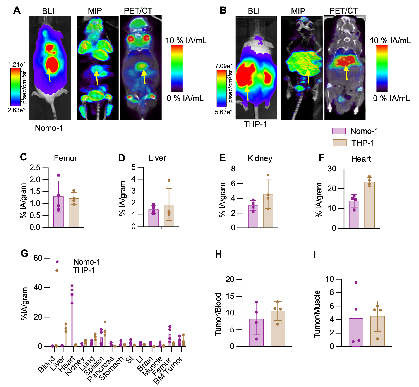

**Figure S13 [^18^F]-FDG PET/CT imaging can detect sites of disease in AML disseminated models, with lower tumor to background**. BLI, MIP and μPET/CT fusion images of [^18^F]-FDG imaging in A) Nomo-1-Luc model (n=4) B) THP-1-Luc model (n=3). ROI drawn on μPET/CT images at C) femur D) liver E) kidney F) Heart pool G) Ex vivo biodistribution indicating %IA/gram accumulation of [^18^F]-FDG in Nomo-1-Luc model and THP-1-Luc model at 1 hour post injection in different organs. H) Tumor/Blood ratio and I) Tumor/Muscle ratio.

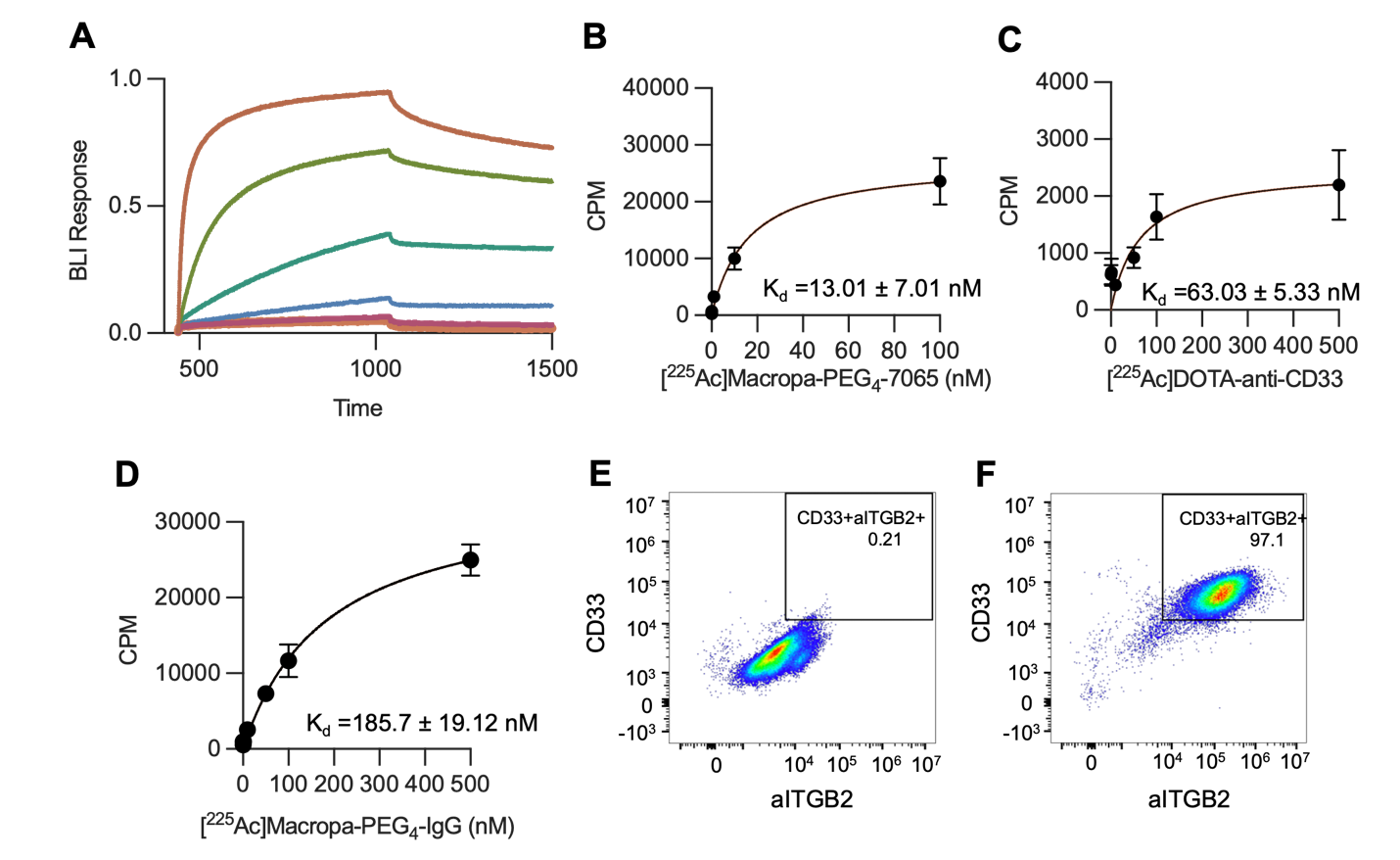

100 nM

20 nM

4 nM

0.8 nM

0.16 nM

0.032 nM

(minutes)

**Figure S14** Representative biolayer interferometry association and dissociation curves between streptavidin-immobilized CD33 recombinant protein and soluble lintuzumab anti-CD33 antibody. Top curve represents binding to lintuzumab ligand at 100 nM, with subsequent curves representing 5-fold dilutions to a minimum of 0.032 nM. Curves have had background absorbance subtracted and baseline aligned utilizing reference sensors and wells.

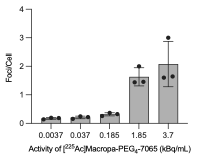

**Figure S15** Immunofluorescence staining for DNA damage (H2AX foci) on the Nomo-1 cells after treatment with different concentrations of [^225^Ac]Macropa-PEG_4_-7065 ranging from 0.0037 kBq/mL to 3.7 kBq/mL for 96 hours.

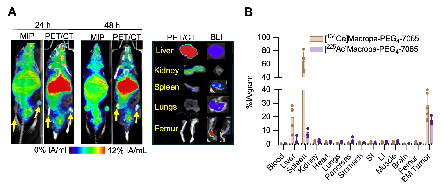

**Figure S16 [^134^Ce]Macropa-PEG_4_-7065 demonstrated high uptake in Nomo-1 disseminated model and served as surrogate to monitor the uptake of [^225^Ac]Macropa-PEG_4_-7065** A) MIP and μPET/CT fusion images of [^134^Ce]Macropa-PEG_4_-7065 in Nomo-1-Luc bearing mouse at 24 h and 148 h post injection. PET/CT and BLI images of the organs injected with [^134^Ce]Macropa-PEG_4_-7065 at 168 h post injection B) Ex vivo biodistribution study comparing the %IA/gram accumulation of [^134^Ce]Macropa-PEG_4_-7065 and [^225^Ac]Macropa-PEG_4_-7065 in all organs at 168 h post injection

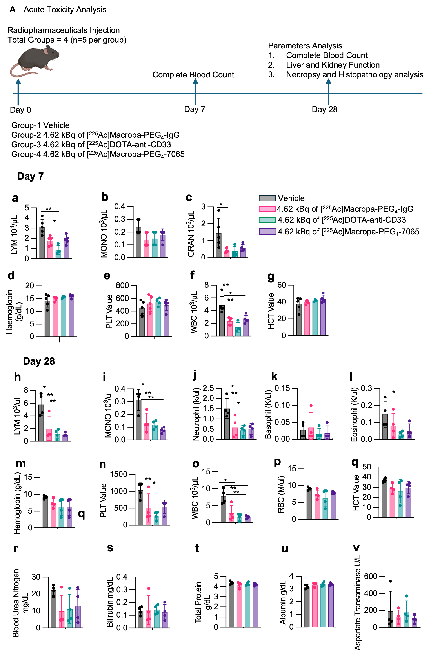

**Figure S17** **Acute toxicity analysis of [^225^Ac]Macropa-PEG_4_-7065 other ^225^Ac conjugates in healthy mice** A) Schematic illustration of Acute toxicity analysis in healthy black C57BJ/6J. Comparison of a) Lymphocytes b) Monocytes c) Granulocytes d) Hemoglobin e) platelets f) WBC g) HCT value among Vehicle, [^225^Ac]Macropa-PEG_4_-7065, [^225^Ac]Macropa-PEG_4_-IgG, [^225^Ac]DOTA-anti-CD33 groups at day 7 post administration of radiopharmaceuticals or vehicle control. Comparison of h) Lymphocytes i) Monocyte j) Neutrophil k) Basophil l) Eosinophil m) Hemoglobin n) Platelet value o) WBC p) RBC q) HCT value r) Blood Urea Nitrogen s) Bilirubin t) total protein u) albumin v) Aspartate transaminase among Vehicle, [^225^Ac]Macropa-PEG_4_-7065, [^225^Ac]Macropa-PEG_4_-IgG, [^225^Ac]DOTA-anti-CD33 groups at 28 days post administration of radiopharmaceuticals or vehicle control.

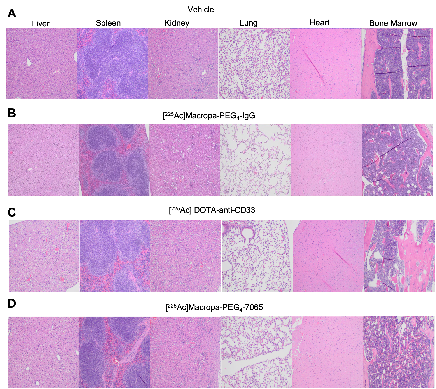

**Figure S18** **Necropsy and Histopathological analysis of liver, spleen, kidney, lung, heart, bone marrow for one of representative mouse in** A) Vehicle B) [^225^Ac]Macropa-PEG_4_-IgG C) [^225^Ac]DOTA-anti-CD33 and D) [^225^Ac]Macropa-PEG_4_-7065 injected cohort.

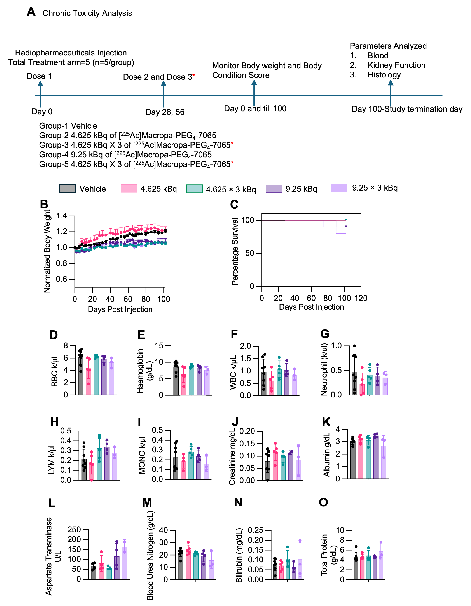

**Figure S19** **Chronic toxicity analysis of [^225^Ac]Macropa-PEG_4_-7065 in healthy mice** **and comparison vehicle group** A) Schematic illustration of Acute toxicity analysis in healthy black C57BJ/6J. B) Body weight plot indicating the measurements in all respective groups C) Kaplan Meier curve demonstrates median time period survival in single as well as fractionated 4.25 kBq and 9.25 kBq dose of [^225^Ac]Macropa-PEG_4_-7065 injected groups respectively as compared to saline. Comparison of D) RBC E) Hemoglobin F) WBC G) Neutrophil H) Lymphocytes I) Monocytes J) Creatinine K) Albumin L) Aspartate transaminase M) Blood Urea Nitrogen N) Bilirubin O) total protein value among vehicle and different doses of [^225^Ac]Macropa-PEG_4_-7065 injected groups.

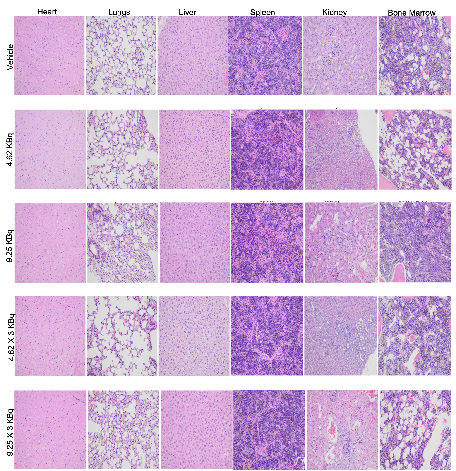

**Figure S20** **Necropsy and** **histopathological analysis of dose-limiting organs** injected with Saline (control group), 4.62 kBq single dose of [^225^Ac]Macropa-PEG_4_-7065, 9.25 kBq single dose of [^225^Ac]Macropa-PEG_4_-7065, 4.62 kBq fractionated dose of [^225^Ac]Macropa-PEG_4_-7065 and 9.25 kBq fractionated dose of [^225^Ac]Macropa-PEG_4_-7065

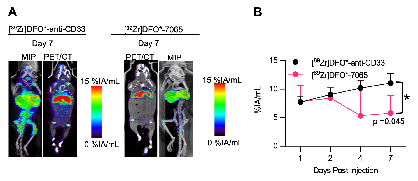

**Figure S21 PET/CT imaging in hCD34+ humanized NOX EXL mice.** A) Fusion PET/CT and MIP images of [^89^Zr]DFO*-anti-CD33 and [^89^Zr]DFO*-7065 at day 1, day 2, day 4 and day 7 post injection B) ROI analysis indicating %IA/gram over a period of 7 days after administration of [^89^Zr]DFO*-CD33 and [^89^Zr]DFO*-7065.

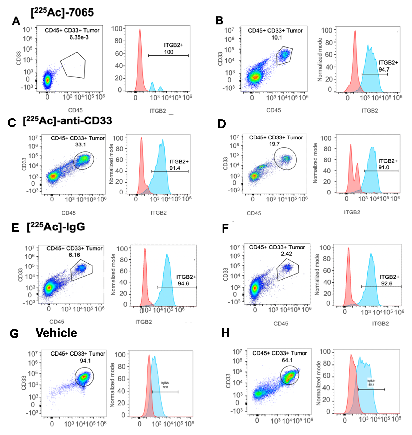

**Figure S22 Flow analysis of relapsed mice from respective therapeutic group in Nomo-1 AML disseminated models.** Flow cytometry analysis indicating CD45+ CD33+ tumor population and aITGB2 expression in bone marrow for other two relapsed mice injected with A), B) 9.25 kBq [^225^Ac]Macropa-PEG_4_-7065. Flow cytometry analysis indicating CD45+ CD33+ tumor population and aITGB2 expression in bone marrow for other two relapsed mice injected with C), D) 9.25 kBq [^225^Ac]DOTA-anti-CD33. Flow cytometry analysis indicating CD45+ CD33+ tumor population and aITGB2 expression in bone marrow for other two relapsed mice injected with E), F) 9.25 kBq [^225^Ac]DOTA-Macropa-PEG_4_-IgG. Flow cytometry analysis indicating CD45+ CD33+ tumor population and aITGB2 expression in bone marrow for other two relapsed G), H) vehicle mice.

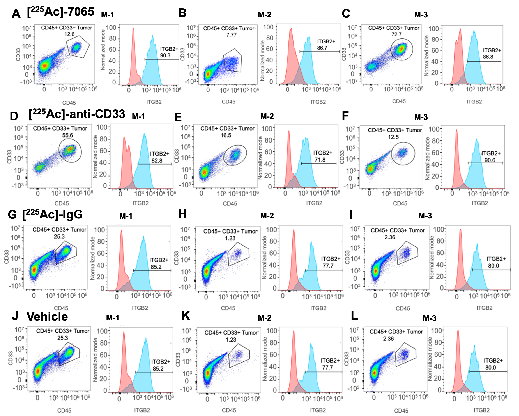
**Figure S23 Flow analysis of relapsed mice from respective therapeutic groups in Nomo-1 AML disseminated models.** Flow cytometry analysis indicating CD45+ CD33+ tumor population and aITGB2 expression in spleen for three relapsed mice injected with A), B), C) 9.25 kBq [^225^Ac]Macropa-PEG_4_-7065. Flow cytometry analysis indicating CD45+ CD33+ tumor population and aITGB2 expression in spleen for three relapsed mice injected with D), E), F) 9.25 kBq [^225^Ac]DOTA-anti-CD33. Flow cytometry analysis indicating CD45+ CD33+ tumor population and aITGB2 expression in spleen for three relapsed mice injected with G), H), I) 9.25 kBq [^225^Ac]DOTA-Macropa-PEG_4_-IgG. Flow cytometry analysis indicating CD45+ CD33+ tumor population and aITGB2 expression in spleen for three relapsed J), K), L) vehicle mice

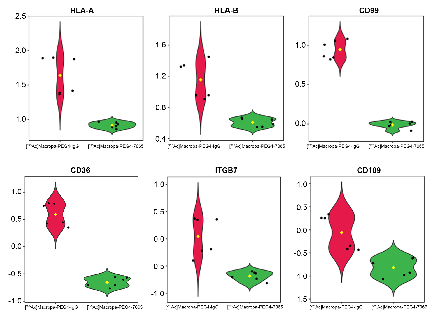

**Figure S24 Specific examples of altered proteins after 7065-based therapy.** Violin plot showing normalized, scaled expression of example dysregulated proteins with *p*-value < 0.05 and foldchange of greater than 2, derived from analysis shown in Figure 7 of main text.

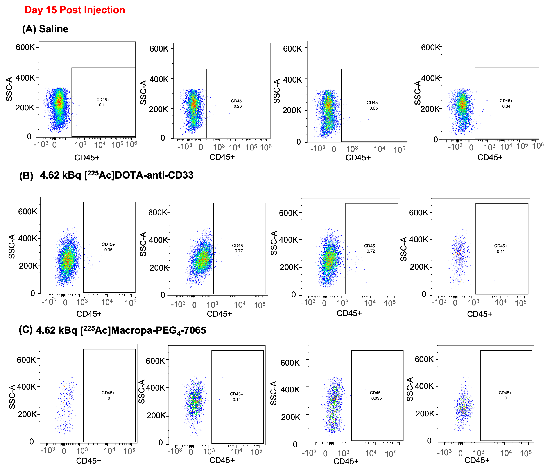

**Figure S25 Flow cytometry analysis for quantification of CD45+ population following treatment in PDX models** at day 15 post injection in blood in A) vehicle B) 4.62 kBq of [^225^Ac]DOTA-anti-CD33 group C) 4.62 kBq of [^225^Ac]Macropa-PEG_4_-7065 injected group.

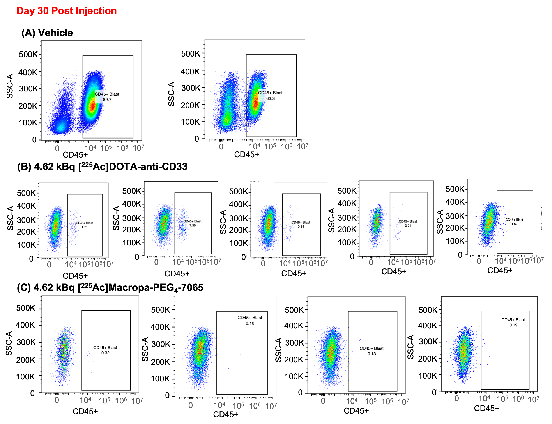

**Figure S26 Flow cytometry analysis for quantification of CD45+ population following treatment in PDX models** at day 30 post injection in blood in A) vehicle B) 4.62 kBq of [^225^Ac]DOTA-anti-CD33 group C) 4.62 kBq of [^225^Ac]Macropa-PEG_4_-7065 injected group.

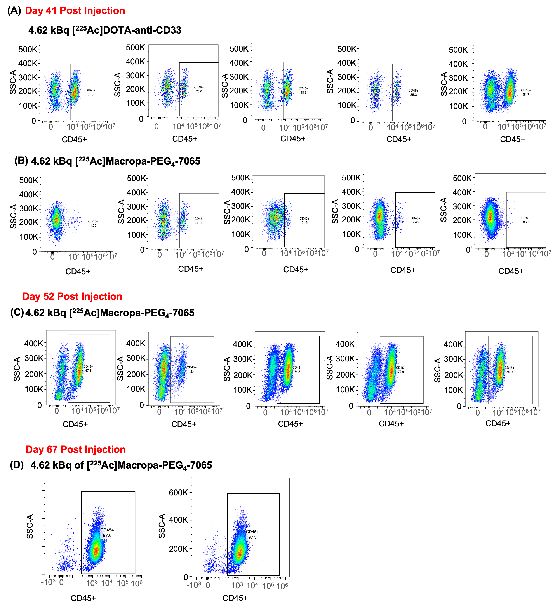

**Figure S27 Flow analysis for quantification of CD45+ population following treatment in PDX models** in blood in A) 4.62 kBq of [^225^Ac]DOTA-anti-CD33 group B) 4.62 kBq of [^225^Ac]Macropa-PEG_4_-7065 injected group at day 41 post injection. Flow analysis for quantification of CD45+ population in blood for [^225^Ac]Macropa-PEG_4_-7065 injected cohort at day C) day 52 and D) day 67 post injection.

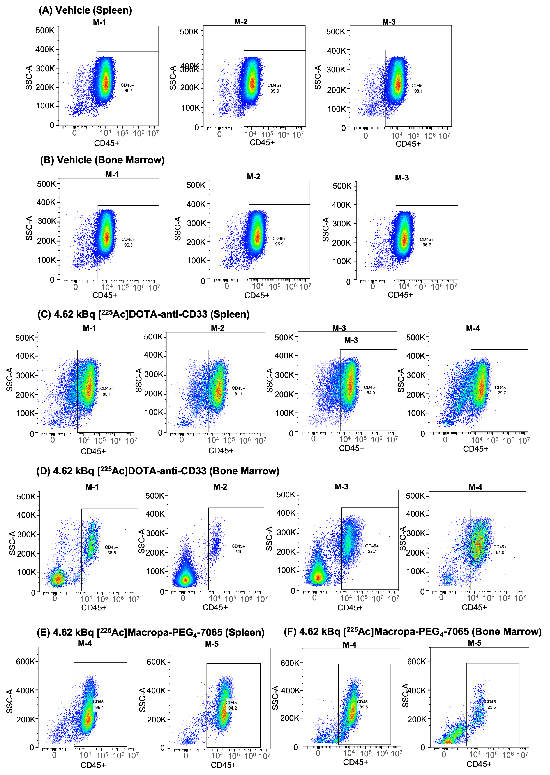

**Figure S28 Flow cytometry analysis for quantification of CD45+ population following treatment in PDX models** in A) spleen B) Bone marrow in vehicle group. Flow analysis for quantification of CD45+ population in C) spleen D) Bone marrow in [^225^Ac]DOTA-anti-CD33 injected cohort. Flow analysis for quantification of CD45+ population in E) spleen F) Bone marrow in [^225^Ac]Macropa-PEG_4_-7065 injected cohort.

**Table S1: Primary Sample Summary Characteristics**

| Total Number of samples | N = 15 |
| --- | --- |
| Samples positive for aITGB2 (MFI fold change > 2) | N = 11 (73.3%) |
| Chromosomal Loss/Gain | N = 9 (60.0%) |
| FLT3 mutation | N = 10 (66.7%) |
| NPM1 mutation | N = 5 (33.3%) |
| DNMT3A mutation | N = 7 (46.7%) |
| IDH mutation | N = 1 (6.7%) |
| TET2 mutation | N = 5 (33.3%) |

**Supplementary Table 1:** Summary characteristics of AML primary samples analyzed for aITGB2 expression. *Abbreviations:* MFI: median fluorescence intensity; FLT3: FMS-like tyrosine kinase 3; NPM1: nucleophosmin; DNMT3A: DNA methyltransferase 3 alpha; IDH: isocitrate dehydrogenase; TET2: Tet methyl cytosine dioxygenase 2

**Table S2: Individual Primary Sample Data**

| **Sample Identifier** | **Fold Change aITGB2 MFI** | **Percent positive aITGB2** | **Sample Type** | **Collection time point** | **Disease blast burden** | **Chromosomal Loss/Gain** | **FLT3 mutation** | **NPM1 mutation** | **DNMT3A mutation** | **IDH mutation** | **TET2** |
| --- | --- | --- | --- | --- | --- | --- | --- | --- | --- | --- | --- |
| 1501 | 1.5 | 24.48 | Leukapheresis | Diagnosis | 100 | ND | ITD, mutation | mutation | ND | ND | ND |
| 1528 | 6.1 | 62.2 | Leukapheresis | Diagnosis | 100 | ND | ITD, mutation | ND | ND | mutation | ND |
| 1520 | 1.8 | 30.6 | Leukapheresis | Diagnosis | 100 | 11q23 (MLL), 11 gain | mutation | ND | mutation | ND | mutation |
| 1400 | 3.0 | 49.61 | Bone Marrow | Diagnosis | 90 | 4 gain, 9 loss | ND | ND | ND | ND | mutation |
| 1367 | 4.1 | 68.1 | Bone Marrow | Diagnosis | 91 | 5p gain, 8 gain, | mutation | ND | ND | ND | ND |
| 1282 | 36.0 | 91.06 | Leukapheresis | Diagnosis | 100 | 6 gain, 8 gain, 5 gain | ND | mutation | mutation | ND | mutation |
| 1475 | 61.9 | 97.7 | Leukapheresis | Diagnosis | 100 | ND | ITD, mutation | ND | mutation | ND | mutation |
| 1493 | 1.5 | 29.83 | Leukapheresis | Diagnosis | 100 | 5 gain, 8 gain, 6 gain, 10 gain, 13 gain | ITD, mutation | mutation | mutation | ND | ND |
| 1513 | 9.2 | 74.3 | Leukapheresis | Diagnosis | 100 | ND | ITD, mutation | mutation | mutation | ND | ND |
| 1536 | 7.9 | 90.7 | Bone Marrow | Diagnosis | 90 | 7 loss, 8 gain, 13 gain | ND | ND | ND | ND | mutation |
| 1646 | 1.3 | 28.9 | Bone Marrow | Diagnosis | 32 | 5q loss, 7q loss, 8 gain, 11q23 (MLL), 2 gain, der(7;21), 12p gain, 13 loss, 16 loss, | ND | ND | ND | ND | ND |
| 1653 | 2.2 | 46.9 | Bone Marrow | Diagnosis | 42 | 8 gain, 20q deletion | ITD, mutation | ND | mutation | ND | ND |
| 1655 | 3.6 | 62 | Bone Marrow | Diagnosis | 72 | ND | ITD, mutation | ND | ND | ND | ND |
| 1658 | 8.6 | 86.6 | Leukapheresis | Diagnosis | 90 | ND | ITD, mutation | mutation | mutation | ND | ND |
| 1680 | 13.6 | 87.4 | Bone Marrow | Diagnosis | 40 | t(1;3), 2q gain, 4q gain, 5q loss, 7q gain, 11q gain, 12 loss, 13 loss, 16q gain, 17p loss | ND | ND | ND | ND | ND |

**Supplementary Table 2:** Individual characteristics of N=15 AML primary samples analyzed for aITGB2 expression. *Abbreviations:* ND: not detected; ITD: internal tandem duplication; MLL: mixed lineage leukemia

**Table S3: Individual Diagnosis/Relapse Primary Sample Data**

| **Sample Identifier** | **Sample Type** | **Collection time point** | **Disease blast burden** | **Time Elapsed from Initial Diagnosis (months)** | **Cytogenetics** | **FLT3 mutation** | **NPM1 mutation** | **DNMT3A mutation** | **IDH mutation** | **TET2** |
| --- | --- | --- | --- | --- | --- | --- | --- | --- | --- | --- |
| 0373 | Bone Marrow | Diagnosis | 80 | 0 | 46,XY,ins(11;9)(q23;p21p24)[20] | ND | ND | ND | ND | ND |
| 0373 | Bone Marrow | Pers/Recur | 95 | 6.5 | 45-46,XY, der(7)t(7;11)(p22;p13), der(11)t(7;11)ins(11;9)(q23;p21p24) [20] | ND | ND | ND | ND | ND |
| 0790 | Leukapheresis | Diagnosis | 100 | 0 | 46XY | ITD, mutation | ND | ND | ND | ND |
| 0790 | Bone Marrow | Pers/Recur | 44-80 | 10.5 | 44-46,XY, t(6;19)(p21.3;p13.3)[4]/46,XY, t(7;9)(p15;q34)[2]/46,XY,del(5)(q22q35)[1]/46,XY,del(16)(q22)[2]/46,XY[11] | ND | ND | ND | ND | ND |
| 0467 | Bone Marrow | Diagnosis | 90 | 0 | 47,XY,+11[4]/48-49,idem,+11,+13[26]. | ND | ND | mutation | mutation | ND |
| 0467 | Leukapheresis | Relapse | 100 | 39.9 | 41-46,XY,dic(7;11)(q11.2;p14),+11[30]. | ND | ND | mutation | mutation | ND |
| 1239 | Bone Marrow | Diagnosis | 23-50 | 0 | 46,XX | mutation | ND | ND | ND | ND |
| 1239 | Peripheral Blood | Pers/Recur | 6.96 | 17.3 | ND | ND | ND | ND | ND | ND |
| 1293 | Bone Marrow | Diagnosis | 50 | 0 | 46,XX,del(20)(q11.2q13.1)[20] | ITD, mutation | ND | mutation | ND | ND |
| 1293 | Bone Marrow | Resid/Recur | 88 | 13.2 | 46,XX,t(1;2)(p34.1;p21),del(20)(q11.2q13.1)[16]/46,XX,t(12;13)(p13;q14),del(20)(q11.2q13.1)[3]/46,XX[1] | ITD, mutation | ND | ND | ND | ND |
| 1377 | Bone Marrow | Diagnosis | 24 | 0 | 46-47,XY,+8 | ND | ND | ND | ND | ND |
| 1377 | Bone Marrow | Pers/Recur | 12 | 6.0 | 43-47,XY,+8[7]/46,XY[3] | ND | ND | ND | ND | ND |
| 1291 | Leukapheresis | Diagnosis | 97 | 0 | 46,XX | ITD, mutation | mutation | mutation | ND | ND |
| 1291 | Peripheral Blood | Relapse | 82 | 35.6 | 46,XX | ITD, mutation | mutation | mutation | ND | ND |

**Supplementary Table 3***:* Individual characteristics of N=14 AML primary samples analyzed for aITGB2 expression collected at diagnosis and again after relapse from venetoclax and azacytidine treatment.

**References**

1. K. Mandal, G. Wicaksono, C. Yu, J. J. Adams, M. R. Hoopmann, W. C. Temple, A. Izgutdina, B. P. Escobar, M. Gorelik, C. H. Ihling, M. A. Nix, A. Naik, W. H. Xie, J. Hübner, L. A. Rollins, S. M. Reid, E. Ramos, C. Kasap, V. Steri, J. A. C. Serrano, F. Salangsang, P. Phojanakong, M. McMillan, V. Gavallos, A. D. Leavitt, A. C. Logan, C. M. Rooney, J. Eyquem, A. Sinz, B. J. Huang, E. Stieglitz, C. C. Smith, R. L. Moritz, S. S. Sidhu, L. Huang, A. P. Wiita, Structural surfaceomics reveals an AML-specific conformation of integrin β2 as a CAR T cellular therapy target. *Nat Cancer* **4**, 1592–1609 (2023).

2. Nix, M. A., Mandal, K., Geng, H., Paranjape, N., Lin, Y. H. T., Rivera, J. M., ... & Wiita, A. P. (2021). Surface proteomics reveals CD72 as a target for in vitro–evolved nanobody-based CAR-T cells in KMT2A/MLL1-rearranged B-ALL. Cancer discovery, **8**, 2032-2049 (2021).

3. S. K. Sharma, A. Chow, S. Monette, D. Vivier, J. Pourat, K. J. Edwards, T. R. Dilling, D. Abdel-Atti, B. M. Zeglis, J. T. Poirier, J. S. Lewis, Fc-mediated Anomalous Biodistribution of Therapeutic Antibodies in Immunodeficient Mouse Models. *Cancer Res* **78**, 1820–1832 (2018).

4. A. Wadhwa, S. Wang, B. Patiño-Escobar, A. P. Bidkar, K. N. Bobba, E. Chan, N. Meher, S. Bidlingmaier, Y. Su, S. Dhrona, H. Geng, V. Sarin, H. F. VanBrocklin, D. M. Wilson, J. He, L. Zhang, V. Steri, S. W. Wong, T. G. Martin, Y. Seo, B. Liu, A. P. Wiita, R. R. Flavell, CD46-Targeted Theranostics for PET and 225Ac-Radiopharmaceutical Therapy of Multiple Myeloma. *Clinical Cancer Research* **30**, 1009–1021 (2024).

5. M. G. Stabin, R. B. Sparks, E. Crowe, OLINDA/EXM: The Second-Generation Personal Computer Software for Internal Dose Assessment in Nuclear Medicine.

6. R. Bruderer, O. M. Bernhardt, T. Gandhi, Y. Xuan, J. Sondermann, M. Schmidt, D. Gomez-Varela, L. Reiter, Optimization of Experimental Parameters in Data-Independent Mass Spectrometry Significantly Increases Depth and Reproducibility of Results. *Mol Cell Proteomics* **16**, 2296–2309 (2017).

7. V. Demichev, C. B. Messner, S. I. Vernardis, K. S. Lilley, M. Ralser, DIA-NN: neural networks and interference correction enable deep proteome coverage in high throughput. *Nat Methods* **17**, 41–44 (2020).

8. A. Barpanda, D. Biswas, A. Verma, S. Parihari, A. Singh, S. Kapoor, C. Kantharia, S. Srivastava, Integrative Proteomic and Pharmacological Analysis of Colon Cancer Reveals the Classical Lipogenic Pathway with Prognostic and Therapeutic Opportunities. *J. Proteome Res.* **22**, 871–884 (2023).
